# Incubator-Free Organoid Culture in a Sealed Recirculatory System

**DOI:** 10.1101/2025.09.03.673593

**Authors:** Yohei Rosen, Kivilcim Doganyigit, Santhosh Arul, Eliot Wachtel, Drew Ehrlich, Vaishnavi Venuturimilli, Matilda Mouzaya, Sophia Oriaku, Bi Ndiforchu, Gregory Kaurala, Sebastian Hernandez, Maryam Moarefian, Hunter Schweiger, Ariana Cisneros, Mojtaba Zeraatkar, Samira Vera-Choqqueccota, Jinghui Geng, Tal Sharf, Sofie Salama, Mohammed A. Mostajo-Radji, Ethan Winkler, David Haussler, Mircea Teodorescu

**Author notes:** These authors contributed equally to this work (Yohei Rosen, Kivilcim Doganyigit). These authors jointly supervised this work (David Haussler, Mircea Teodorescu).

## Abstract

Discovery in human biology is pivoting toward high-dimensional computational analysis of 3D in vitro models, but this progress is limited by reliance on conventional cell culture techniques. Realism and data collection are hindered by the environmental instabilities and accessibility constraints of standard incubators. We introduce an automated, sealed recirculatory system that eliminates these barriers, enabling unconstrained instrument integration and infrastructure-independent scalability. By employing gas-tight sealing, a liquid-phase gas buffer and a non-porous plastic gas exchanger, our technology maintains biological stability without the compromises of open-air vessels. This design eliminates the need for CO**_2_** incubators and prevents the evaporative drift that typically plagues conventional open-culture vessels. Operating on the benchtop outside the cell culture suite, we demonstrate that our system supports continuous, multi-week live imaging of vascular organoids while maintaining metabolic viability, structural fidelity and electrophysiological activity in brain organoids comparable to traditional in-incubator cultures.

**Teaser:** A sealed benchtop device enables data-rich organoid culture beyond the constraints of the cell culture laboratory.

## Introduction

The landscape of biomedical discovery is undergoing a historic transformation. Progress in data handling and machine learning offer exponential growth in the scale of analyses possible for biological experiments. Meanwhile, global regulatory shifts such as the FDA Modernization Act 2.0 [1] now prioritize human-relevant research over traditional animal testing. With a transition to prioritizing so-called next-generation physiological systems [2, 3], in vitro models aiming to closely replicate in vivo tissues, human-derived organoids have moved from a niche interest to a regulatory necessity. But existing laboratory constraints waste the potential of organoids. A primary benefit of organoids over animal models is the ability to observe them freely and continuously instead of relying on invasive endpoint measurements [4, 5]. Time series observations of organoid models can yield detailed data on organogenesis [6, 7], disease pathogenesis [8] and function [9], which would be difficult to obtain in vivo. The utility of organoids will be maximized by techniques that improve their physiological fidelity and increase observability for high dimensional data collection [10].

Organoids face two barriers to achieving this mandate: the logistical difficulty of maximizing data collection and continued shortcomings in physiological realism. Modern data handling capabilities and machine learning models are beginning to allow generalist users to analyze large, complex datasets. [11, 12] But there is a mismatch between these analytical tools and the limited data throughput of most biological experiments. Research is constrained by manual labor and the inherent difficulty of integrating tissue culture with longitudinal observation [10].

The Petri dish, the cell culture suite and the CO2 incubator date back to the 19th century. [13] These technologies were designed with human operators in mind. Over the past century, the equipment has improved but the principles remain fundamentally unchanged, impeding automation. The next generation of cell culture experiments require automation, scaling and integration of advanced sensors [14, 15]. Currently, manual handling and the physical constraints of traditional cell culture suites create a physical barrier to sensor access, and technician-dependent infrastructure limits throughput and temporal resolution of data collection. New approaches to cell culture are needed to remove bottlenecks for AI-driven discovery.

Organoid biology also requires continued technological development to attain the realism necessary to fully replace animal models. Accuracy of modeling is critical to the success of next-generation physiological models for preclinical discovery. Traditional cell culture labware relies on open-air gas exchange, which limits realism by creating a microenvironment that diverges from mammalian physiology, with the tissue-air interaction of an open wound rather than the natural interaction of healthy internal tissue with the body. In conventional cell culture devices, this wound-like air-liquid interface is used to regulate tissue oxygen and pH by CO_2_-supplemented air equilibration (Fig. 1e). But open-air culture triggers a cascade of biochemical failures. Evaporation is not merely a logistical nuisance; it is directly cytotoxic [16–18] and has been shown to fundamentally alter cortical network activity [19]. Unlike the air-liquid interface in cell culture, most cells in the body rely on interstitial fluid that is regulated and isolated from the external environment. In vivo, tissues exist in closed compartments (Fig. 1a(i)), with interstitial fluid microenvironments [20] (Fig. 1a(ii)) tightly regulated by microvasculature [21–25] (Fig. 1a(iii)). This fluid environment is further regulated by the homeostatic functions of organs like the kidneys and liver [26], with additional regulation from the lungs [27] (Fig. 1a(vi)). To mimic this, homeostasis ought to be performed in a similarly closed environment.

**Fig. 1.**
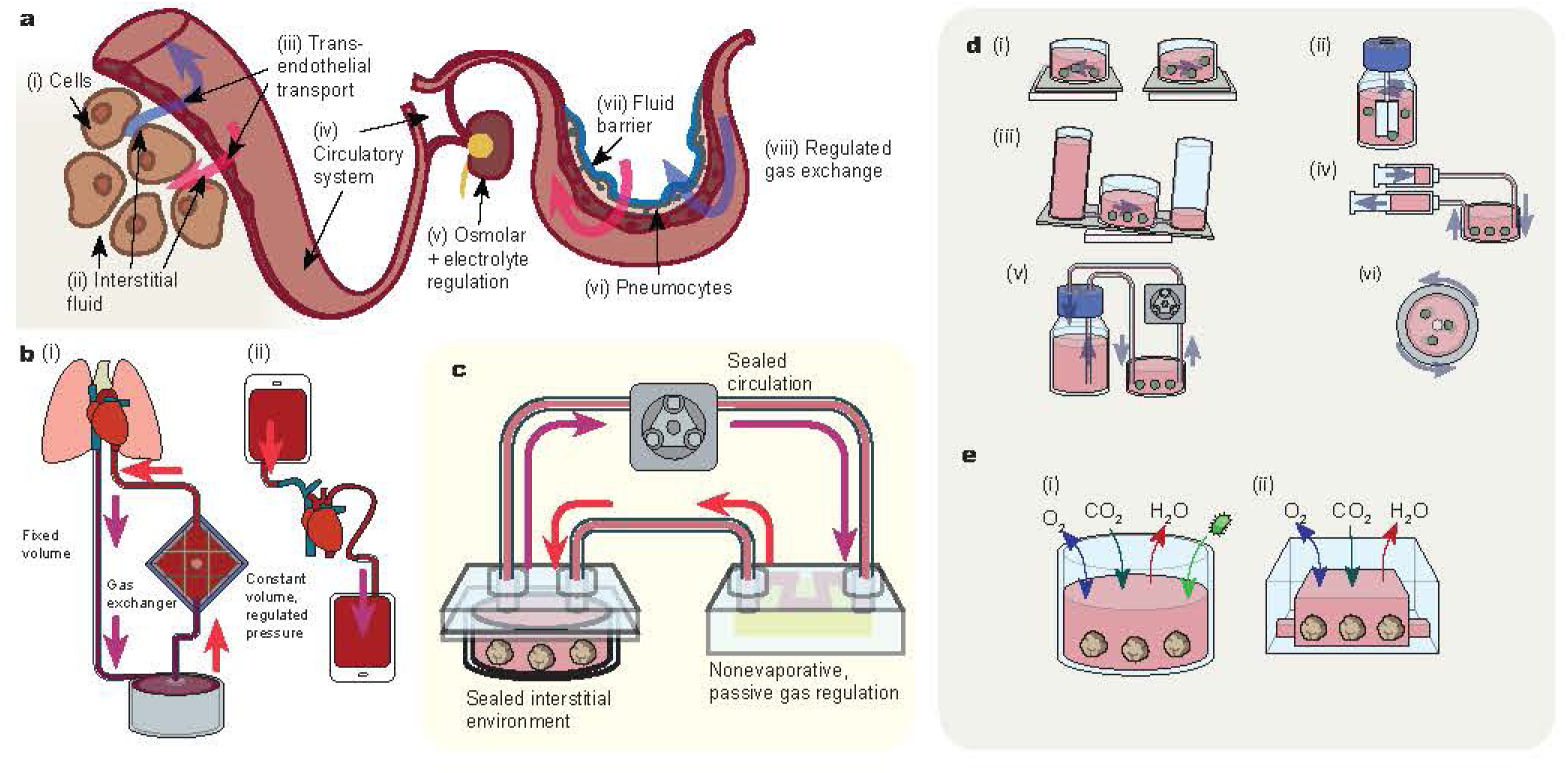
Design principles for incubator-free culture a, Schematic of mammalian homeostasis: Nonepithelial tissues (i) are nourished by diffusion and advection of interstitial fluids (ii), with composition regulated by transendothelial transport (iii). Circulating plasma composition (iv) is further regulated by organs, e.g. in renal osmolar and electrolyte control (v). Plasma gas content is regulated in the alveoli by ventilation-perfusion matching (vi-viii). **b,** Homeostasis in intensive care medicine. (i): ECMO exchanges gas outside of the body in an air-free sealed circuit. (ii): Simultaneous exchange transfusion replaces blood without significantly altering circulating volume or pressure. **c,** Our approach to circulatory organoid culture includes a sealed cell culture media environment, a sealed circulatory media loop and a non-evaporative gas exchanger for passive gas regulation. **d,** Existing approaches to circulation/agitation in cell culture: (i): inertial shaker plates, (ii): stirred bioreactors, (iii): rocker well gravity circulation, (iv): syringe or air displacement circulation, (v): peristaltic pump circulation, (vi): rotating wall bioreactors. **e,** Schematic of gas exchange, evaporation and contamination in (i) air-liquid interface systems such as in well plates or gas-balanced reservoirs and in (ii) silicone devices such as PDMS microfluidics.

Organoids also require improvements in nutritive transport. In the body, microvasculature perfuses the interstitial space, exchanging fluids and gases through pressure gradients and active transport (Fig. 1a) [23, 24, 28]. In contrast, organoids lack perfused vasculature and are often too thick for effective diffusion, limiting metabolic exchange [29]. This impairs their physiological realism. Spheroids commonly exhibit a necrotic core due to insufficient nutrient supply [29]. Some incubator-based organoid culture strategies partially address diffusion challenges by enhancing surface advection, which renews concentration gradients and may promote permeation through porous tissues [29, 30]. This enables the outer layer of organoids to experience fluid transport similar to perivascular interstitial spaces, improving on static diffusion-limited conditions. But forced advection combined with an air liquid interface increases evaporative toxicity [31].

We assert that the ideal system for organoid culture in the era of data and automation should have three features distinguishing it from current methods. It should allow straightforward access for continuous observation, whether with high frequency microscopy or with more specialized instruments. It should allow scaling unhindered by infrastructure limitations. It should maximize its physiological realism, in order to provide the most accurate environment for biomedical discovery.

In this work, we report a cell culture system with all these features, breaking away from cell culture labware convention dating back to Petri. [13] By replacing the physiological drawbacks of the air-liquid interface [16–19] with a hermetically sealed, recirculatory flow architecture, we establish a microenvironment that mirrors internal mammalian physiology far more accurately than open-well systems. By creating an accessible benchtop platform, we provide for unimpeded continuous imaging and chemical sensor monitoring of tissue development and state progression for both scientific study and feedback control of tissue state trajectory. The increased biological realism, scalability and realtime controllability of our system has the potential to elevate organoid research from boutique to mainstream in biomedical research.

The core innovation in our system is to replace the air-liquid interface with flat sheet polymethylpentene gas exchanger. Using an improved version of the “liquid-breathing” aqueous buffer approach from the Futai group [32], we eliminate evaporation and eliminate the need for gas feedback control. This design also totally seals the device, freeing it from secondary containment and other environmental constraints which necessitate the expensive and inflexible standard cell culture suite infrastructure for conventional incubator culture. Our sealed system can run human tissue experiments in any lab, while conventional incubator culture requires a biosafety level 2 laboratory. Our fluidically sealed gas exchanger also allows a fixed-volume circulatory loop with single-actuator media exchange through volume displacement (Fig 3e), yielding simplified fluidic handling and a miniaturized system. Critically, our design creates a self-enclosed device that is much more compact than an incubator (Fig. S19), easily accessible to instrumentation (Figs. 2d, 4b, Fig. S20) and isolated from its environment for asepsis (a major issue in organoid culture [33]) and containment.

**Fig. 2.**
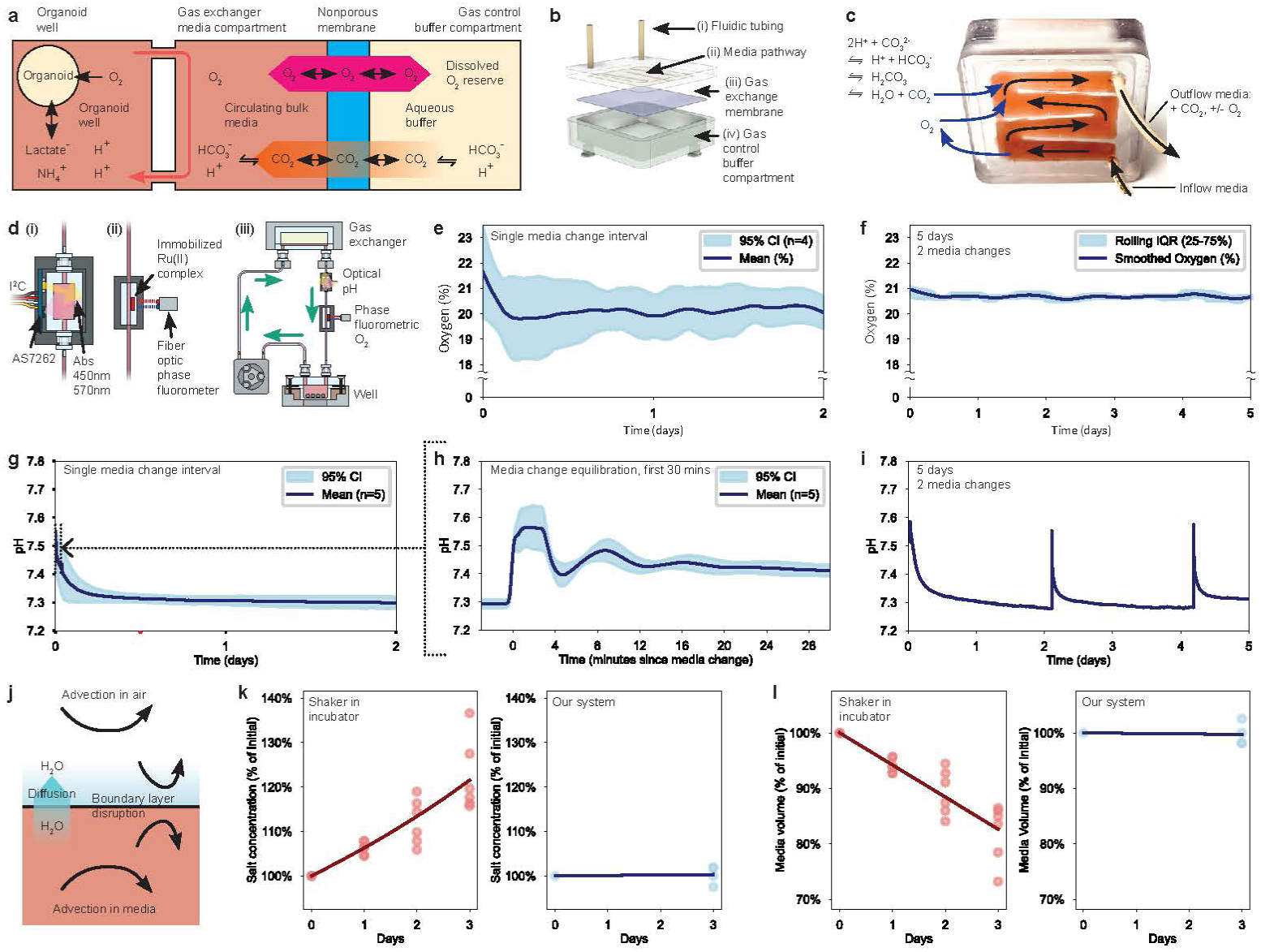
Gas transport and homeostasis in incubator-free cell culture. a, In a gas control buffer compartment, CO_2_ is generated by an aqueous buffer. O_2_ and CO_2_ diffuse freely through non-porous polymethylpentene. CO_2_ is circulated to the organoid media environment, interconverting with carbonic acid to maintain physiological pH despite the production of ammonium, lactic acid, and other acids from cellular metabolism. **b,c,** Design of planar gas exchanger. Media flows into a 200 *µ*m deep serpentine pathway (ii) through fluidic tubing (i). The media passes over a polymethylpentene membrane (iii), exchanging gases with an aqueous buffer (iv) by diffusion. **d,** Design (i,ii) and arrangement (iii) of pH and O_2_ optical sensors. **e,f,** Phase fluorometric series measurement of O_2_ shows oxygen homeostasis following equilibration from initial media hyperoxia. **g,** Time series measurement of pH shows equilibration to a homeostatic range of pH 7.30–7.34, with an initial equilibration from hypocarbic, alkalotic fresh media via combined mixing and diffusion. Dotted region shows extent of time plotted in **h**. **h,** First 30 minutes of pH equilibration after media change (subset of **g**), dominated by fluid mixing with residual media. **i,** 5 day measurement of media pH, spanning 2 media changes. **j,** Schematic of factors contributing to free water loss by evaporation and advection in systems using an air-liquid interface and especially in agitated culture systems: boundary layer disturbance and bulk transport refreshing concentration gradients. **k,** Relative salt concentration change for shaker in incubator culture and in our system. Trend fit to [*Na*^+^](*t*)*/*[*Na*^+^]_0_ (%) = 100*/*(1 *− At*), for incubator *A* = 0.059 days*^−^*^1^ (95% CI 0.051–0.057), for our system *A* = 0.001 days*^−^*^1^ (95% CI -0.007–0.009). For difference, p = 0.0003 by *t*-test. **l,** Calculated free water volume change for shaker in incubator culture and in our system. Trend fit to *V ol*(*t*)*/V ol*_0_ (%) = 100 *− Bt*, for incubator *B* = 5.8 %/day (95% CI 5.0–6.6); for our system *B* = 0.093 %/day (95% CI -0.74–0.93). In **k,l,** Curves fit by least squares regression. In **e,g,h,** data are shown as Savitzky-Golay smoothed pointwise mean of replicate measurements and 95% CIs calculated by pointwise *t* -tests. In **f**, smoothed data are a Savitzky-Golay filtered 10^4^ measurement rolling window and rolling IQR over the same window.

**Fig. 3.**
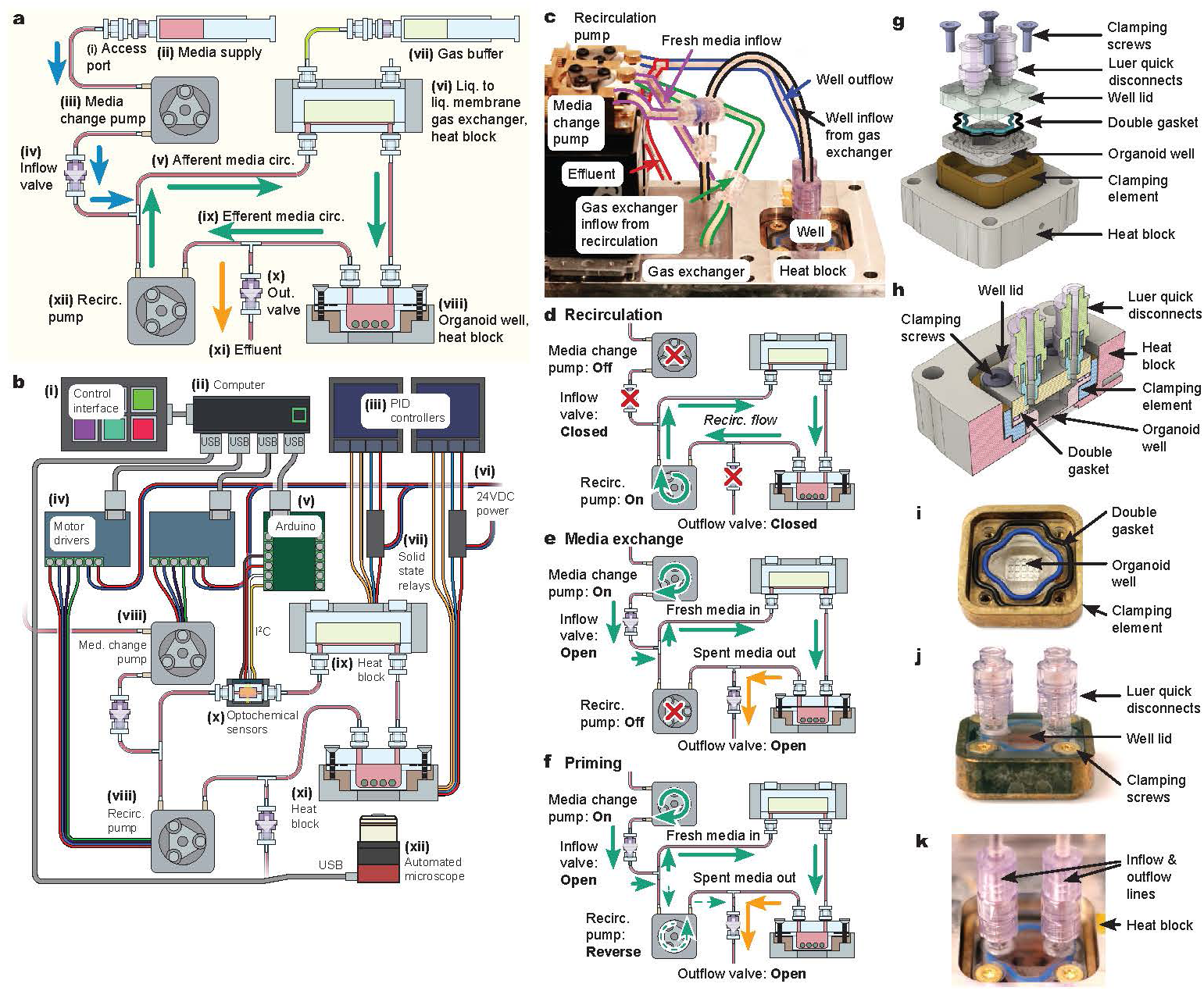
System design and function a, A refrigerated syringe used as media reservoir (ii) connects to fill port (i). A peristaltic pump (iii) actuates media changes by injection across a check valve (iv) attached to the recirculation loop via tee junction. Media enters the afferent side (v) of the recirculation circuit and passes through a liquid to liquid PMP membrane gas exchanger (vi). Gas buffer exchange is via syringe (vii). Media flows through an organoid well (viii) into the efferent side (ix) of the recirculation loop. A peristaltic pump (xii) recirculates efferent media. When media is injected by media change pump (iii), this also opens outflow check valve (x) shunting efferent media to an effluent line (xi). **b,** An Elgato Streamdeck (i) or command line provides a human interface to a computer (ii). It communicates over USB with motor drivers (iv) which drive peristaltic pumps for fluidic actuation (viii). Standalone PID controllers (iii) control heat block (ix, xi) temperature via solid state relays (vii). The system is powered by 24V DC current (vi). The computer acquires automated microscope images using a bespoke USB microscope (xii). Optionally, a colorimetric pH sensor (x), coupled via Arduino (v) or a phase fluorometric oxygen sensor are included inline and monitored over USB. **c,** Photograph of assembled apparatus as used for vascular organoid culture. **d,** Recirculation is effectuated via recirculation pump with the system sealed to the external environment via closed inflow and outflow valves. **e,** Media change is effectuated by inflow via media change pump and shunting to outflow by stopping the recirculation pump. **f,** Priming, or full clearance of the system is effectuated similarly to media change by with the recirculation pump running in reverse to clear the portion of the loop downstream of the outflow valve. **g-k,** Design of organoid well. Organoids sit on the bottom of a polycarbonate well. Screws seal the well by clamping the well lid, double gasket and well against a clamping brace. The metal clamping brace inserts into the heat block and conducts heat to the organoid well.

**Fig. 4.**
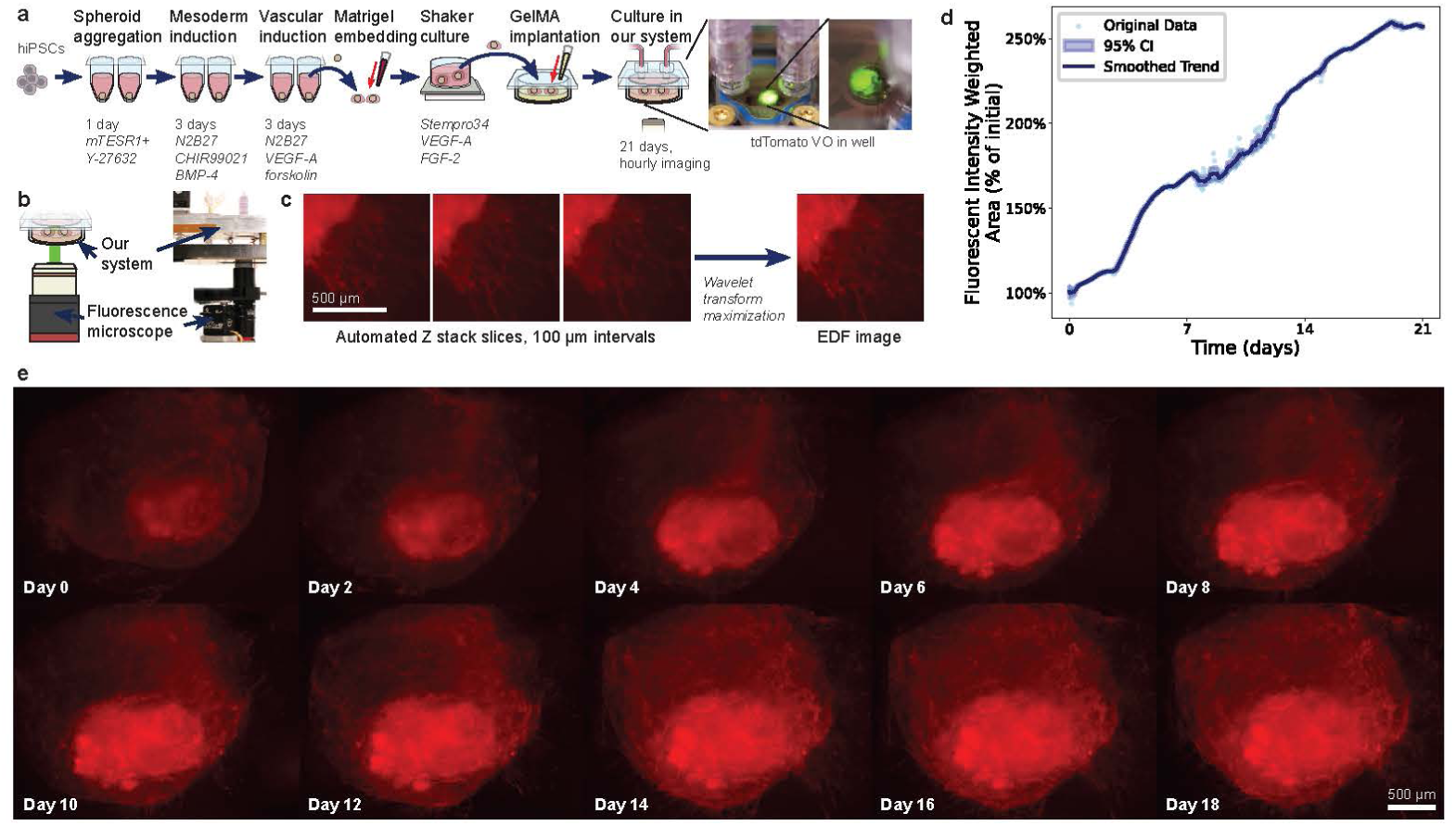
Vascular organoid culture and long-term epifluorescence imaging. a, Human vascular organoids were prepared from WTC11-tdTomato hiPSCs according to a version of the Wimmer protocol. They were embedded in Matrigel, maintained on shaker until use, loaded in our system in a GelMA matrix and cultured for 21 days with automated fluorescence microscopy. **b,** Microscopy apparatus for live cell imaging of our system incubator. **c,** Method of generating extended depth of field (EDF) composites via pixelwise complex wavelet transform maximization, shown using Z-stack images of a subregion of vascular outgrowths. **d,** Growth curve calculated by summing fluorescent signal over the area of organoid images, using average per-pixel intensity between Z planes, smoothed with a 10-image rolling window. CI shows pointwise 95% CI calculated by *t*-test across this rolling window. **e,** Vascular organoid, showing inner globular zone and outer vascular network zone. Representative subset of ten imaging timepoints, from 21 days of hourly images, demonstrating timepoint to timepoint imaging consistency. Images are EDF composites and were gamma corrected (*γ* = 0.8) in batch for visual appearance. Scale bar indicates 500 µm.

## Results

### System design for incubator-free organoid culture

A central drawback to traditional cell culture labware is its reliance on open-air gas exchange. This complicates both realism and access to cultures. We made our design decisions to address these limitations. Long term culture is generally performed in CO2 incubators, which are sealed from the room to maintain CO_2_ and humidity levels and containment. Sealing interferes with instrument access, confining cultures in cramped, rigid and environmentally harsh [16–19] surroundings. Microscope access is sometimes improved by short term use of stagetop incubators, but these generally have insufficient gas and humidity homeostasis for long term culture and are also inflexible closed chambers. Air-contacting cultures are also vulnerable to contamination. [33] Aseptic handling and environmental isolation are needed, an additional barrier to access. Liquid handling in these systems is also open-air, requiring airspace isolation for antisepsis and droplet containment. Primary airspace isolation requires laminar flow hoods, and if biosafety control is needed, secondary containment with room-level airspace isolation. With these physical restrictions in place, limited laboratory space restricts replication, equipment inflexibility restricts the nature of experiments, and need for fluid handling to happen in isolation cabinets limits the complexity of equipment integrations for observation and automation.

We have designed our cell culture system to eliminate the incubator and air-liquid interface, improving accessibility to instrumentation, improving fidelity to the mammalian tissue microenvironment and completely sealing organoid culture. (Figs. 1c, 2a) Our general design mimics the self-contained environment of mammalian tissues, where cells are in closed compartments isolated from outside air, with gas homeostasis through membrane diffusion and fluid circulation (Fig. 1a). Our apparatus consists of a hermetically sealed organoid well with optical windows for imaging, an air-free sealed recirculation loop with automated media exchange and a nonporous membrane gas exchanger. With this design, isolation is not required surrounding the cell culture vessel.

Achieving such a design depends on a gas exchange method to maintain media gas homeostasis other than an air-liquid interface. In intensive care medicine, when the lungs cannot be used, extracorporeal membrane oxygenation (ECMO) heart-lung machines (Fig. 1b(i)) achieve gas exchange through nonporous membrane gas exchangers, isolating blood from room air. We adopt this approach in eliminating the air-liquid interface present in conventional incubators. To simplify the system and add a secondary barrier against evaporation, our gas control medium is an aqueous carbonate-buffer. [32] This achieves pH homeostasis (through buffering) and osmolar homeostasis (by preventing evaporation) eliminating the need for complex feedback control of either parameter.

### Comparison to existing systems for improving media transport in organoid culture

To improve over the diffusion limitations of static culture, organoid culture is often performed with forced advection of media. The two primary methods for forced advection are media agitation devices and circulation devices [29]. Shaker culture (Fig. 1d(i)) [34] is commonly used for inertial agitation. Rocker fluidic devices [35–38] (Fig. 1d(iii)) use gravity and custom geometries for more structured advection. These use variants of traditional well plates, with all the disadvantages that come with them. Roller [39] (Fig. 1d(vi)) and stirred impeller bioreactors [4] (Fig. 1d(ii)) also promote bulk advection while minimizing impact forces through flotation. While agitation is simple, it causes random displacement, complicating longitudinal studies like imaging or electrophysiology as well as mechanical trauma. Non-agitatory methods are crucial for data rich in vitro biology. Agitation at the air-liquid interface also worsens the physiological divergence of tissue cultures from real organs by accelerating evaporation [31, 40].

The principal alternative to agitation is media circulation through vessels containing statically positioned organoids. To prevent organoid movement and trauma, media circulation devices flow media through open [30, 41] or closed [42, 43] tissue culture chambers for advection. Methods include peristaltic pump recirculation (Fig. 1d(v)), seen in [42, 43] and pneumatic [44, 45] or syringe pumping [30] (Fig. 1d(iv)). Circulation is promising for ease of instrument integration, but gas exchange in reported systems is still either open air or through poor vapor barriers.

Circulation and agitation partially alleviate the diffusion-limited nutrition of static organoid culture [29, 30] but not other deviations from physiological realism. Improvements in advection in air-contacting systems are complicated by worsened evaporation such that improved nutritive flow is often at the cost of less realistic osmolyte homeostasis. While advection enhances transport at the organoid surface, it disrupts the air-liquid interface [40, 46], potentially increasing evaporation compared to static cell culture dishes [31] (Fig2j). Air contact is obvious in well plate cultures [34, 36–38], but most circulatory devices, including closed wells, have similar air exposure from static [43], agitated or bubbled [47] air-contacting reservoirs used to provide gas exchange, and thus face the same evaporative challenges. Even PDMS [16, 48] and silicone tubing [49, 50] gas exchange designs encounter comparable evaporation issues [16] (Fig. 1e(ii)). Due to their high water vapor permeability, PDMS microfluidics fluidics often impair biological function [16] or suffer circulatory failures from evaporative salt precipitation or bubbling.

In contrast, our system achieves circulation without creating a more evaporative air-liquid interface. In this manner, we are able to improve media transport as compared to static plate culture, without agitatory displacement of organoids and without osmotic derangement interface.

### Design of a liquid-breathing gas exchanger for O_2_ and CO_2_ regulation

For gas control, we used a two-compartment, air-free system that regulates culture gases via pH and oxygen buffering in a fluidically isolated chamber. We improved on the Futai group’s method [32] using a thermoplastic membrane to avoid PDMS’s biochemical disadvantages [51] and high water vapor permeability [16] while improving manufacturability [52].

We use the biocompatible polymer polymethylpentene (PMP) [53] for gas exchange due to its high gas permeability. Our exchanger (Fig. 2a-c) features a media path fluidically isolated from a bicarbonate buffer chamber by a nonporous PMP membrane, providing CO_2_ and O_2_ homeostasis. (Fig. 2e-i)

### A nonporous gas exchanger maintains oxygen and pH homeostasis

Oxygen stability during cortical organoid culture was assessed using oxygen phase fluorometry (Fig. 2d) inline with the gas exchanger. Oxygen concentrations remained stable within the normal cell culture range. Excluding the initial 3 hours post-media change, mean oxygen readings (N=5) were 19.6–20.4%, with the pointwise 95% confidence band spanning the interval of 18.2–21.5% O_2_ (Fig. 2e). Similar stability was observed over a 5-day culture period (Fig. 2f).

We tracked pH during cortical organoid culture using ratiometric absorbance spectroscopy of media phenol red using a digital color sensor affixed to a flowthrough cuvette (Fig2d) according to the calibration curve shown in Fig. S9. We observed rapid equilibration from hypocarbic fresh media after media changes (Fig. 2g,h). Median time for to equilibrate to below pH 7.4 was 0.6 hours, IQR 0.1–0.9 hours. Excluding the initial 3 hours after media change, pointwise median pH (N=5) was stable in the range of 7.30–7.34 with the pointwise 95% confidence band spanning the interval of 7.28–7.39 (Fig. 2g). These results were further reflected in a 5 day culture period (Fig. 2i). pH homeostasis as a measure of CO_2_ containment also demonstrates gas-tight sealing since continued efflux via an incomplete seal would raise pH above 8.

### Liquid- and gas-sealed organoid culture achieves osmotic homeostasis

Osmotic homeostasis was evaluated through sodium concentration changes, allowing calculation of electrolyte concentration disturbance and volume loss. Data was gathered over three-day trials with 2 mL starting volume. A reciprocal linear fit for electrolyte concentration and linear fit for volume loss showed significant improvement over shaker in incubator culture (*t* -test, *p* = 0.0003, Fig. 2k,l). In shaker culture, 114 µL excess media free water was lost per day from a 2 mL starting volume (95% CI 87–141 µL/day), while our system showed negligible loss (95% CI -1.5–2.0 µL/day) (Fig. 2l). We observed high evaporation in incubator with other plate formats and significant evaporation inhomogeneity (Figs. S2–S7) consistent with [17].

### Design of an air- and expansion reservoir-free automated recirculation culture loop

Our design goals were sealing, reliability and actuator simplicity. As discussed above, we required a system which achieves media circulation in a sealed system. Fundamentally, our system is a peristaltic pump-driven recirculatory loop, however we were able to minimize the circulating media volume and adopt a novel media change strategy due to the benefit of an air-free, constant volume sealed apparatus.

Most in vitro organoid systems require variable enclosed volumes for sequential removal and addition of fluid during media exchange. Conventional setups (e.g., flasks, wells) use the culture chamber as a variable-volume reservoir, while other systems employ remote expansion reservoirs [42, 43, 54]. With a variable volume, automating media changes necessitates multiple coordinated actuators, which increases system size, costs and failure points. [55, 56]. These factors hinder the ability to scale cheaply or reliably, which is critical for the large scale automation required for AI driven laboratories. Expansion reservoirs also create excess dead volume, diluting endogenous signals and added reagents, potentially altering cell behavior and reducing measurement sensitivity. Actuators in open-air systems may inadvertently aspirate air bubbles, disrupting gas concentrations, precipitating solutes, and causing turbulence, blockage, or inconsistent mixing that affects cells, sensors, or flow. Coordination can be challenging due to actuator errors or volume inconsistencies and may necessitate feedback control to manage variability in fluid handling steps [57].

Our system, which has no air-liquid interface, consists of sealed rigid-walled elements with a fixed internal volume that enables media exchange through displacement by injected media. This approach is simpler than the multi-actuator methods used in other circulatory culture designs. In medicine, simultaneous arteriovenous exchange transfusion (Fig. 1b(ii)) enables fluid exchange without disrupting circulating volume. In this work, we apply these methods to organoid culture.

The system comprises:

- A fixed volume media recirculation loop
- An air-free organoid well (Fig. 3a.viii)
- A gas exchanger (Fig. 3a.vi)
- A recirculation pump (Fig. 3a.xii)
- An inflow pump entering the loop via tee junction (Fig. 3a.iii)
- An outflow line attached via a tee junction
- Inflow and outflow check valves preventing backflow (Fig. 3a.iv,x)

All elements are sealed to fluids and gases; fluidic access is achieved through commercial intravenous line aseptic access ports.

Recirculation (Fig 3d) uses a peristaltic pump returning efferent media from the well to the gas exchanger before reentering the well. Absent media injection, the inflow and outflow check valves remain closed, sealing the system.

To exchange media (Fig. 3e), the media change pump injects fresh media while the stalled recirculation pump occludes recirculation. The injected media displaces spent media into the outflow line, which can be coupled to further analytical instruments (Fig. S20). Recirculation is halted while maintaining uninterrupted flow through the organoid well. Unique to our system, it is alternatively possible to perform continuous trickle media change without need to coordinate inflow and outflow actuators (Fig. S18).

Priming the system (Figure 3f) involves running the recirculation pump in reverse while otherwise following the media change protocol. The pump shunts spent media to outflow and clears its own dead volume, enabling complete media replacements.

We designed the system with computer-based controllers (Fig. 3b) which minimize disturbances to organoids and enable integration with automated sensing modules (Fig. 2d, 4b), in line with other recent work on automation of organoid culture [57].

### Sealed well design and antisepsis

The organoid well is a sealed module with a screw-clamped lid and self-sealing ports for circulatory system integration (Fig. 3g,j,k). It features a double gasket: a compressible fluorosilicone inner O-ring for fluidic isolation and an outer Aflas FEPM O-ring for gas blocking (Fig. 3g,i).

A sealed design prevents gas leakage and eliminates the need for clean storage of the culture apparatus, allowing culture on a non-sanitized workbench. This design avoids the access and ergonomic challenges of using a biosafety cabinet or laminar flow hood and incubator, as well as the difficulty of maintaining asepsis in a fluidic system. Despite comprising many modules, there are no open junctions for microbial ingress; only the well and media injection port require initial aseptic loading. Once sealed and connected (Fig 3k), they no longer need a clean environment.

### Our apparatus achieves long-term benchtop culture of human vascular organoids with high-frequency time-series fluorescence imaging

To demonstrate our system’s utility for long-term organoid culture and dynamic morphological observation, we cultured human vascular organoids (see Methods) for 21 days with high-frequency live cell epifluorescence imaging. Matrigel-embedded WTC11-tdTomato human iPSC-derived vascular organoids (Wimmer protocol [58]) were cultured in our system on gelatin methacrylamide (Fig. 4a). A miniaturized fluorescence microscope acquired images through the well bottom (Fig. 4b). Fluorescent intensity weighted area measurements showed our system supports robust growth over multiple weeks and accommodates undisturbed high-frequency imaging during media changes (Fig. 4d). Complete fluidic automation ensured consistent imaging. A representative subset of time series images is shown in (Fig. 4e) without between-timepoint alignment.

### Characterization of murine cortical organoids cultured on benchtop in our system

For further biological validation, we cultured murine embryonic stem cell-derived cortical organoids (COs) [59] in our system (see Methods). Cohorts of 12-16 COs were cultured for 5.5 days and compared to matched cohorts in well plates under shaker [34] and static incubator conditions (Fig. 5a). Four independent cohorts were used, aged 12 to 33 days at experimental day 0.

**Fig. 5.**
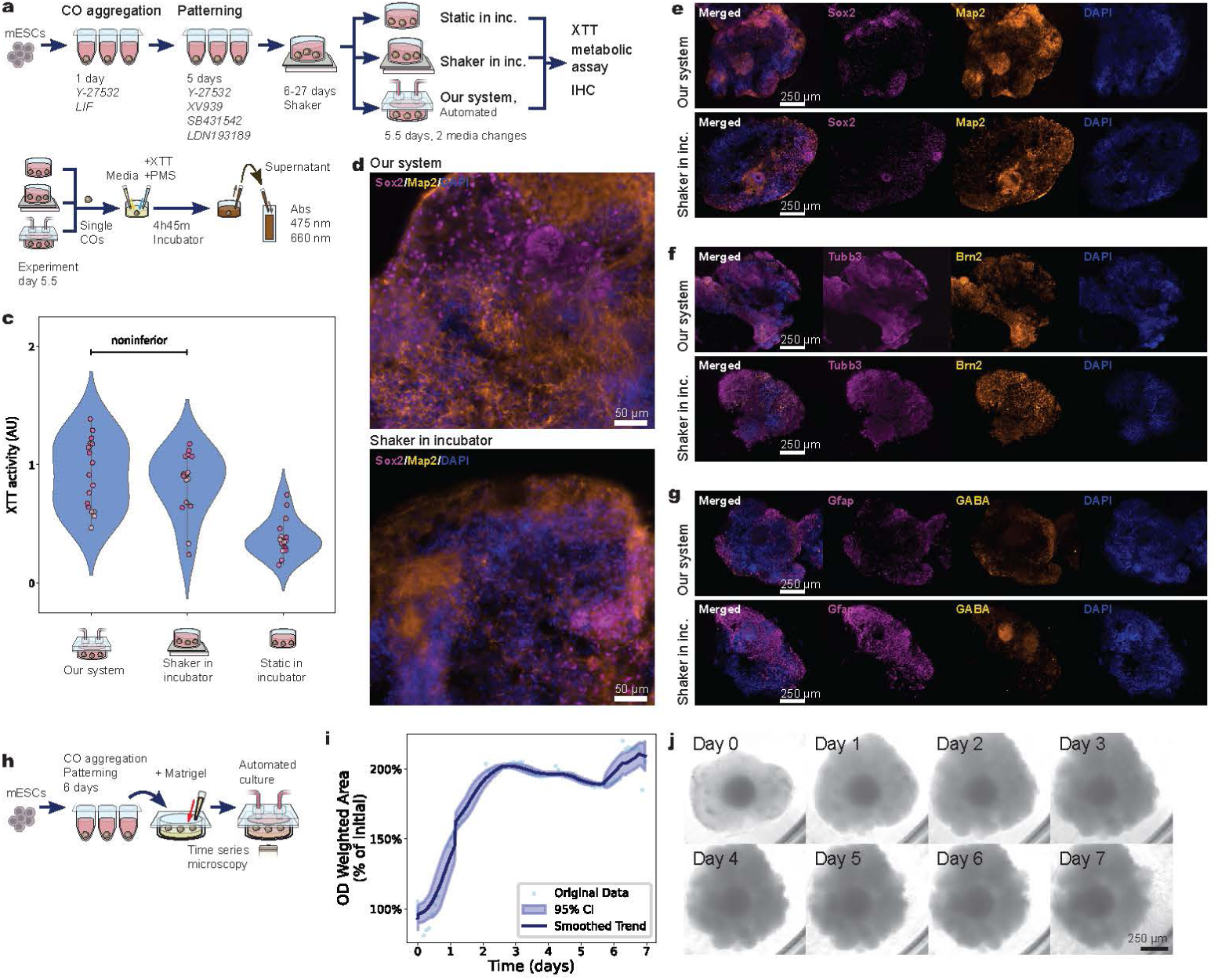
Cortical organoid culture and characterization a, Protocol for cortical organoid culture for XTT and IHC assays. **b,** Protocol for XTT NAD(P)H dependent oxidoreductase activity assay. **c,** XTT metabolic activity from formazan absorbance (AU). Scatterplot colors reflect grouping by experimental cohort. Noninferiority calculated by maximum likelihood estimation mixed effects analysis (controlling for experimental cohort) via pairwise *t*-tests with noninferiority margin Δ = 10% and 95% confidence intervals. 95% CI of relative XTT activity of our system was 92–130% better than shaker in incubator culture, 159–326% better than static incubator culture. **d–g,** Representative fluorescence immunohistochemistry of 40 µm vibratome sections from 32 day old mouse dorsal cortical organoids. Organoids were cultured either in our sealed incubator-free perfusion system (top rows) or on shaker in a conventional incubator (bottom rows). **d,** Merged images of cortical organoid subregions stained for Sox2 (neural progenitors) and Map2 (neurons), with nuclear counterstaining with DAPI. Scale bar: 50 µm. **e,** Full organoid sections stained for Sox2, Map2, and DAPI, and corresponding merged images. Scale bar: 250 µm. **f,** Sections were stained for Tubb3 (neuronal microtubules), Brn2 (deep-layer neurons) and DAPI, and corresponding merged images. Scale bar: 250 µm. **g,** Sections were stained for Gfap (astrocytes), GABA (inhibitory neurons) and DAPI, and corresponding merged images. Scale bar: 250 µm. **h,** Protocol for cortical organoid culture for growth tracking assay. **i,** Growth curve calculated from live cell imaging by summing total optical density across organoid area. **j,** Representative subset of live imaging timepoints demonstrating cortical organoid growth in our system. Images processed by extended depth of field fusion, cropped and brightness scaled in batch.

### Metabolic viability of murine cortical organoids

Metabolic viability of COs was compared via mitochondrial NAD(P)H oxidoreductive metabolism using an XTT assay. 4–5 COs per condition were randomly selected and formazan absorbance was measured (Fig. 5b). Data were analyzed with a linear mixed effects model, treating experimental cohorts as random effects. Pairwise *t*-tests indicated noninferiority (Δ = 10%) of our culture apparatus compared to both shaker and non-shaker in-incubator cultures. 95% CIs for relative XTT metabolic activity showed our system was 92–130% better than shaker in incubator culture and 159–326% better than static incubator culture (Fig. 5c).

### Live cell imaging of murine cortical organoids

We used live-cell microscopy of Day 6 COs cultured in our system for one week to track growth (Fig. 5h). Following the method of [60], we cultured with Matrigel to visualize growth phenotypes less evident in growth factor-free media. We observed increasing CO cross-sectional area (Fig. 5j) and aggregate optical density (Fig. 5i).

### Immunohistochemistry of murine cortical organoids

In order to validate architectural and lineage fidelity of COs grown in our system against those grown using an orbital shaker in an incubator (shaker in incubator) [34, 59], we performed immunohistochemistry (IHC) using dorsal cortical organoid relevant [59] markers. The pictured data (Fig. 5d-g) show day 32 COs after 5.5 days culture.

Map2 (microtubule-associated protein 2) supports mature dendritic architecture [61], while Sox2 marks neural stem and progenitor cells [62, 63]. Stainings for both markers were comparable between our system and shaker in incubator cultures, with rosette-like structures present in both (Fig 5d,e), consistent with intact progenitor organization and neuronal maturation. Additional replicates shown in Fig. S1.

Tubb3 (*β*-III-tubulin), a neuron-specific cytoskeletal component essential for axonal extension and an early neuronal identity marker [64], was strongly expressed in organoids from our system and shaker in incubator culture, particularly in outer zones, suggesting neuronal lineage commitment (Figure 5f). Similarly, Brn2, a transcription factor essential for cortical fate specification and the progenitor-to-neuron transition [65–67] showed similar intensity and patterning across both methods of culture, suggesting comparable regulation of early cortical neurogenesis (Figure 5f).

Astrocytic specification was assessed using the glial marker Gfap, with both organoids from our system and shaker in incubator culture revealing Gfap-positive peripheral and central domains (Figure 5g). Dorsal identity in both conditions was confirmed by detection of GABA (*γ*-aminobutyric acid), an inhibitory neurotransmitter, which is expected to be sparsely present at day 32; results aligned with this expectation in both methods of culture (Fig 5g).

### Electrophysiology of murine cortical organoids

To validate functional development, we cultured day 27 dorsal forebrain COs in our system for 5.5 days and tested them against shaker cultured COs (Fig. 6a). We looked for coherent electrophysiological network activity, expected in mature dorsal forebrain COs[68] using Maxwell Biosystems MaxOne high-density microelectrode arrays. Following established analysis workflows [59, 69, 70], we identified putative firing neuronal units (Fig. 6b) and computed spike-time tiling coefficients (STTCs) [71], a measure of synchrony between pairs (Fig. 6c). COs cultured in our system and cultured on shaker in the incubator before MEA mounting exhibited comparable network synchrony. Electrophysiology measurements were taken at Day 41, at which point both groups displayed modest correlated activity with median STTC values of 0.083 (IQR 0.001–0.193) and 0.042 (IQR 0.009–0.108) respectively. A portion of neuronal pairs surpassed the 0.3 synchrony threshold (10.2% of pairs in our system; 6.7% in shaker), while strong correlations (*>*0.5) were rare (2.1% and 2.2%).

**Fig. 6.**
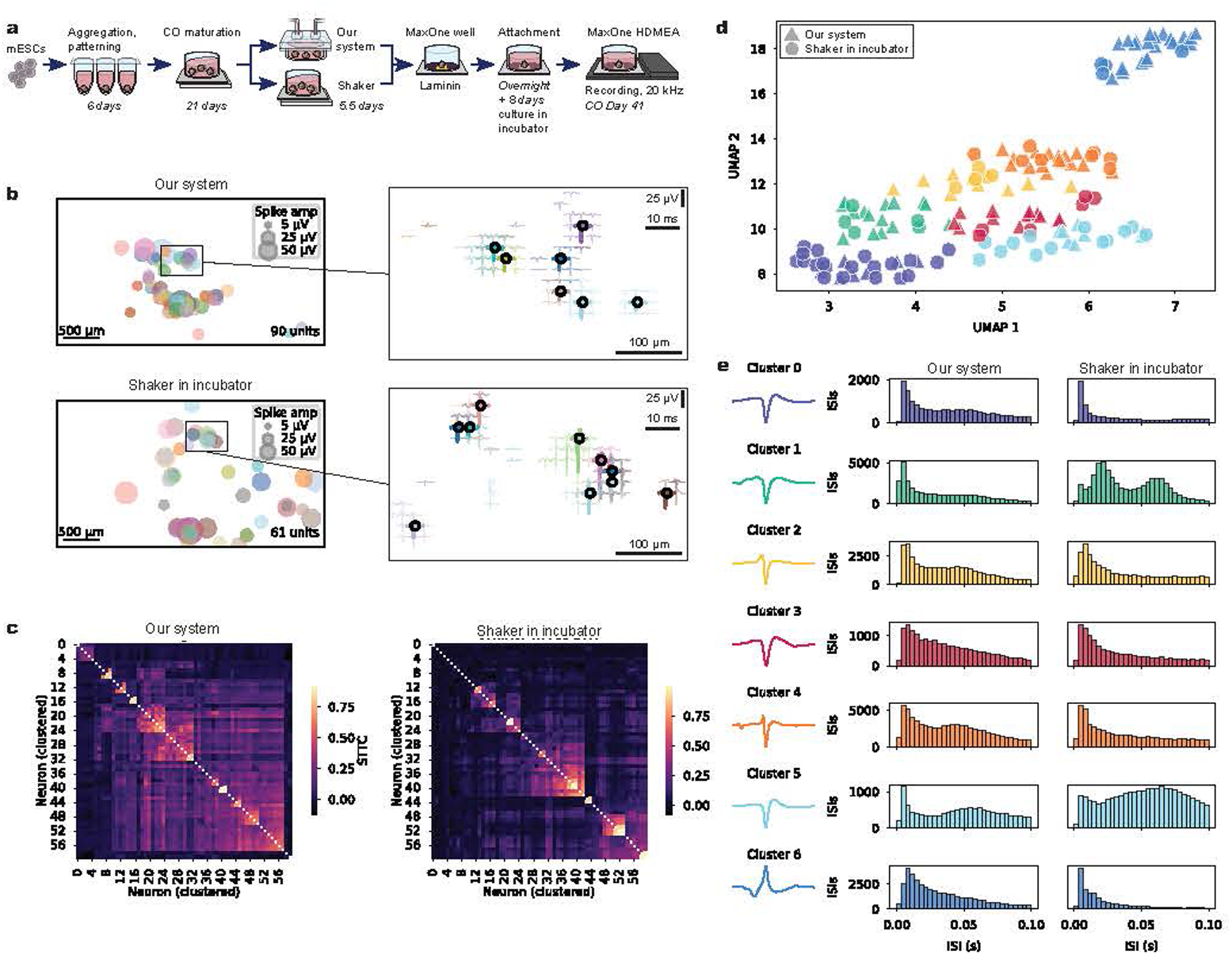
High-density electrophysiological characterization of cerebral organoids a, Schematic of experimental workflow. mESCs were aggregated and patterned into dorsal forebrain cortical organoids (COs) for 6 days, followed by 21 days of suspension maturation. COs were then transferred to either our incubator-free system (*Our System*) or maintained on shaker in a conventional incubator. After 5.5 days, (approx. day 33 of organoid maturity) one organoid per group was attached to a high-density CMOS microelectrode array (Maxwell Biosystems MaxOne HD-MEA; 26,400 electrodes, 17.5 µm pitch) coated with laminin and cultured for 8 days to adapt to the CMOS array plate. Electrophysiological recordings were acquired at a sampling rate of 20 kHz on approx. day 41 of organoid maturity. **b,** Spatial maps of single-unit extracellular spike amplitudes recorded across the 1020 most active electrode sites in each condition. Each bubble indicates the location of a spike-sorted putative unit; bubble size denotes mean binned spike amplitude (5 µV, 25 µV, 50 µV). Spike sorting identified 90 units in the CO from our system and 61 in the the CO grown in incubator on shaker. Insets (right) show overlaid spike waveform footprints from units within the boxed regions. Waveform amplitude and time scale bars: 25 µV (vertical) and 10 ms (horizontal). Electrode position scale bar: 100 µm. **c,** Clustered heatmaps of spike time tiling coefficients (STTCs) computed for the 60 most active units per condition (Δ*t* = 10 s). Hierarchical clustering of average linkage was used to reorganize the STTC matrix such that functionally similar neurons are grouped together. Warmer colors indicate higher synchrony. **d,** Two-dimensional UMAP embedding of per-unit average spike waveforms, colored by cluster identity (*k*-means, *k* = 7). Triangle markers indicate units from our system and circles indicate shaker in incubator culture. Clusters were computed on pooled units across conditions to assess similarity and divergence in waveform space. **e,** Cluster average spike waveforms (left) and inter-spike interval (ISI) histograms (right) for each cluster. Waveforms are color-coded to match panel d. Waveform plots are scaled to normalize maximum absolute amplitudes for comparison of shape while minimizing amplitude bias. ISI histograms, scaled cluster-wise, are shown separately for each condition to illustrate cluster-specific firing patterns and bursting behavior.

We used waveform clustering on pooled data from both conditions to compare cluster assignments between conditions as a measure of waveform consistency. Units from both experimental conditions were identified across all clusters, suggesting comparable organization and diversity of waveform morphologies (Fig. 6d,e). Within each cluster, interspike interval (ISI) histograms revealed varied firing regimes; some clusters exhibited burst-like activity with short intervals, while others showed broader distributions consistent with tonic or irregular spiking (Fig. 6e). This diversity suggests that both culture conditions support a range of electrophysiological phenotypes rather than biasing development toward a single firing regime.

## Discussion

Organoid culture is limited in scale by labor intensiveness and poor access for observation and automation and physiologically by tradeoffs between nutrition and osmotic homeostasis. Organoids provide a potential method to study organ development, function and pathology in high detail, but current methods restrict integration with the instrumentation needed to capture their richness. We have developed a modular apparatus for recirculatory organoid culture that enhances practicality and biological accuracy. Its buffered liquid-–liquid gas exchanger enables controller-free gas homeostasis and improved osmolarity control. The fixed-volume, air-free circulation loop simplifies fluidic design (improving cost and reliability for automation), reduces labor and minimizes sources of variability. Integrated optical chemical sensors and microscopy provide data-rich continuous monitoring of organoid cultures. By eliminating reliance on open incubator systems, we have created a compact, aseptic, evaporation-free automated benchtop culture platform. Our tests with vascular and cortical organoids prove that our system supports growth, viability and differentiation comparable to incubator-based shaker culture while improving osmotic homeostasis and observational capabilities.

The benefits of this apparatus will extend beyond the current study. Its evaporation control is especially valuable for tissue models sensitive to ionic fluctuations [19], such as electrode-integrated brain organoids (including for organoid computing), cardiac and muscle models and transport epithelial models including kidney and mucosal organoids. The sealed, passively gas-controlled design is suitable for studies involving altered gas environments like hypoxia, important for disease modeling. Complete fluidic sealing prevents contamination and isolates toxic or biohazardous samples in a simple benchtop system without requiring conventional laboratory containment. Microbial contamination and cell line cross-contamination are major pain points in academic and industry cell culture; sealed devices prevent both by isolation.

By eliminating the need for large equipment like incubators and biosafety cabinets, organoid culture is freed from the confines of the cell culture laboratory. This enables colocation with diverse analytical instruments (Fig. S20). Sensitive electrode-based sensors require electrical isolation and high-sensitivity live cell microscopes need space and cooling that conflict with incubator culture. Optical detection devices, such as infrared and Raman spectrometers, phase fluorometers (as used in this study), custom laser equipment on optical benches and refractometric or interferometric instruments are also often difficult to integrate into conventional cell culture due to vibration sensitivity, isolation requirements or bulk. Similarly, large monolithic platforms for conditioned media analyses including extracellular vesicle analyzers, realtime mass spectrometers, online flow cytometers and integrated immunoassay systems are commonly used in commercial bioreactor settings but are difficult to integrate into conventional setups. The trend of academic core facilities for central instrumentation further complicates incubator culture integration with analytical instruments. Our system will facilitate greater integration of organoid culture with advanced laboratory instrumentation enabling extensive data collection from tissue models.

Freeing organoid culture from the cell culture suite and the incubator also unlocks unbounded scalability. The system’s compact size and simplicity with minimal actuators and minimal need for controllers naturally lead to scalability in parallel automated devices (Fig. S19). Parallelism will be restricted by total laboratory space, rather than by the limits of existing human-centric cell culture infrastructure. Our system provides a potential foundation for large-scale, automated biology producing high-dimensional data. Such a system will accelerate the development of rapid biomedical discovery in AI-driven laboratories.

## Resource availability

- **Materials availability.** This study did not generate new unique materials/reagents. Computer aided design files for CNC milled and 3D printed device components are available upon request.
- **Data and code availability.** Data and code underlying this paper’s analyses as well as system control scripts are available upon request.

## Funding and Acknowledgements

This work was supported by the Schmidt Futures Foundation SF 857 (S.S., D.H. and M.T.), the National Human Genome Research Institute (NHGRI) under Award number RM1HG011543 (S.S, D.H. and M.T.), by the National Science Foundation (NSF) under awards number 2134955 (S.S, D.H. and M.T.) and 2034037 (M.T.), by the California Institute for Regenerative Medicine (CIRM) awards DISC4-16285 (S.S, M.A.M.-R. and M.T.) and DISC4-16337 (M.A.M.-R.), by the University of California Office of the President award M25PR9045 (S.S., M.T. and M.A.M.-R.), by the National Institute of Mental Health (NIMH) award U24MH132628 (M.A.M.-R. and D.H.) and National Institute of Neurological Disorders and Stroke award U24NS146314 (M.A.M.-R. and D.H.) and NIMH award R01MH120295 (S.S) and the University of California Santa Cruz Center for Information Technology Research in the Interest of Society and the Banatao Institute Interdisciplinary and Innovation Program (I2P) (M.A.M.-R). Y.R. and M.Z. were supported by CIRM postdoctoral fellowships under Award number EDUC4-12759 and the Institute for the Biology of Stem Cells (IBSC) at UC Santa Cruz. Y.R. was supported by the Peggy and Jack Baskin fellowship and UCSC Baskin School of Engineering fellowship programs. Maryam Moarefian was supported by an National Institutes of Health under award number K12GM139185 and the IBSC at UC Santa Cruz. A.C was supported by a T32 fellowship under NHGRI award number T32HG012344. H.S. was partially supported by the NSF Graduate Research Fellowship Program. S.H. received support from the UC Doctoral Diversity Initiative (DDI-UCSC-IBSC). S.V.-C. was partially supported by the Graduate Pedagogy Fellowship from the University of California Santa Cruz Teaching and Learning Center. B.N. was supported by a UCSC CORE fellowship. The technology underlying this work was also supported by an UCSC Innovation Catalyst Grant.

We thank Ryan Fenimore for assistance with assays and contributions to pH sensing software. We thank Jake Revino and John Minnick for assistance with alternative engineering prototypes which were ultimately not used in this work and Isabel Cline for sharing IHC techniques. We thank Ravipa Losakul for discussions regarding integration with instruments. Benjamin Abrams, UCSC Life Sciences Microscopy Center, RRID: SCR 021135 for technical support. Electrophysiology data was hosted on a pipeline built on the National Research Platform (NRP), supported by the National Science Foundation under award numbers CNS-1730158, ACI-1540112, ACI-1541349, and OAC-1826967, as well as by the University of California Office of the President and the UC San Diego California Institute for Telecommunications and Information Technology/Qualcomm Institute.

## Author contributions

Y.R. conceived of and engineered the devices and methods described, assembled the devices and control systems, wrote control and analysis software, conceived of and performed experiments, oversaw all work and authored the paper. K.D. provided feedback contributing to iterative redesign of devices and methods, helped manufacture and assemble parts, conceived of experiments, performed and directed experiments, analyzed data, wrote code for analysis pipelines and authored subsections of the paper. S.A. prepared experimental samples, contributed feedback on engineering and contributed technical assistance for vascular organoid work. Eliot Wachtel contributed to sensor engineering. D.E. contributed to microscopy engineering. V.V., B.N., S.O., Matilda Mouzaya, Maryam Moarefian and A.C. provided experimental samples. G.K. contributed technical and analysis assistance for experimental assays. S.H. provided experimental samples and contributed feedback with respect to protocols. H.S. contributed software tools used in electrophysiology. S.V. assisted with experiments. M.Z. assisted with transparent well manufacturing. J.G. contributed to software tools used in electrophysiology. T.S. provided experimental resources. S.S. advised on experimental methods and provided resources. M.A.M.-R. advised on immunohistochemistry and provided experimental resources. Ethan Winkler provided experimental resources. M.T. and D.H. provided supervision and advised on engineering, experimental methods, data reporting and writing.

## Declaration of generative AI and AI-assisted technologies

During the preparation of this work the authors used ChatGPT to edit text for tone and concision. GitHub CoPilot was used to assist with python scripting in data processing and visualization workflows. After using these tools, the authors reviewed and edited the content as needed and take full responsibility for the content of the published article.

## Supplemental information

- Materials and Methods.
- Supplemental Figures S1-S21, supplemental tables S1-S9.

## Methods

### Device manufacturing materials

Machining stock, gaskets, tooling and fasteners were sourced from McMaster Carr unless otherwise noted. Polymethylpentene was sourced from CS Hyde. Adhesives were sourced from RS Hughes.

### Design and manufacturing of components

3D models were designed in Autodesk Inventor Professional 2023. Steel, brass and aluminum computer numerical control (CNC) milling and metal 3D printing fabrications for custom jigs, heat blocks and fixtures were manufactured by PCBWay’s manufacturing service. CNC milling of plastics was performed in-house on Bantam Tools or Carbide3D Nomad mills with CNC toolpaths generated in Autodesk Fusion using milling tools sourced from Harvey Tool. Ultraviolet light curing used a Loctite CL15 light source. CNC milled parts were deburred using an S-blade and scalpel, rinsed with isopropyl alcohol and air-dryed before further assembly. To remove cutting agents, metal parts were washed sequentially with dish soap (Dawn), water, and isopropyl alcohol, then air-dried prior to use.

### Cell culture well manufacturing

Cell culture wells (Fig. 3g-k) were manufactured from polycarbonate by CNC milling with reverse milling achieved using a custom aluminium jig. For general purpose wells, well bottoms were polished with a 3/16” polycrystalline diamond router bit (LMT Onsrud 75-110). For experiments requiring fluorescence imaging, the bottom of the well was manufactured from a separate piece of polycarbonate and affixed with Loctite 4311. For experiments requiring brightfield imaging (Fig. 5h-j), wells were 3D printed on an Asiga Ultra 32 printer using Detax Freeprint Ortho resin. Bill of materials in Table S1.

### Cell culture well lid manufacturing

Well lids were CNC machined from polycarbonate and 1/4”-28 UNF fluidic ports by thread milling. Lids for brightfield imaging were overlaid with integrated light diffusers constructed from CNC milled 1/8” polypropylene. (Fig. S11)

Female luer lock ports were inserted into the threaded lid ports, sealed with Loctite 4311. (Fig. S12) Self-sealing swabbable luer lock ports (Qosina) were installed on top of these. Bill of materials in Table S2. A double gasket (Fig. 3g,i), consisting of an outer 1/16” round profile size 20 Aflas perfluoroelastomer O-ring (McMaster 5240T211) and an inner 1/16” round profile size 13 fluorosilicone O-ring (McMaster 8333T13), was placed in the gasket glands of the well bottom. (Fig. S10) Sealing of the well lid against the well was achieved using custom brass M4 *×* 0.7 mm 14 mm countersunk flat head Torx drive screws custom manufactured by Shenzhen Shi Shi Tong Metal Products (Fig. 3g-j). Clamping pressure for gasket sealing was achieved by a CNC milled brass clamping element (Fig. 3g-i) with four M4 threaded holes to accept the clamping screws. The clamping element was designed to mate on its inner surface with the bottom of the cell culture well and on its outer surface with the interior of the well heat block (Fig. 3h,k) in order to conduct heat to the cell culture well. Brass hardware was chosen for its passive antiseptic properties.

### Gas exchanger manufacturing

The media side of the gas exchanger (Fig. 2b(ii)) was milled from polycarbonate, featuring 0.2 *×* 10 mm cross section serpentine channels matching the cross-sectional area of 1/16” inner diameter fluidic tubing. Pharmed BPT tubing was bonded directly into 3.5 mm ports using Loctite 7701 and Loctite 4311. A stepped brim was milled around the serpentine channel to enclose the serpentine channel for bonding the polymethylpentene gas exchange membrane. A nonporous 0.003” polymethylpentene film (Fig. 2b(iii)), cut to size by hand, was pretreated with Loctite 7701 polyolefin primer.

The brim and channel dividers of the serpentine channel were coated with a thin, continuous layer of Loctite 4311. The polymethylpentene film was affixed to the adhesive and crosslinked by UV transillumination. After priming the obverse side of the film with Loctite 7701, the perimeter was again sealed to the polycarbonate brim with a second overlaying layer of UV-cured Loctite 4311 as a sandwiching coating.

The buffer reservoir side of the gas exchanger (Fig. 2b(iv)) was milled from 1/2” polycarbonate. Openings milled into the backside of the reservoirs were tapped to 1/4”-28 UNF threads as access ports. The gas exchanger was attached to the prepared media exchange side and polymethylpentene film assembly using Loctite 4011. The perimeter was overcoated with Loctite 4311 cured with UV light. A 3 mm *×* 6.35 mm stirbar was inserted inside the reservoir. Luer ports were constructed by bonding 1/4”-28 UNF thread to female luer adapters into the threaded holes and sealing using Loctite 4311. Gas exchangers were quality controlled by an air leakdown test at 2 psi. Bill of materials in Table S3.

### Peristaltic pump manufacturing

Peristaltic pumps were constructed around a NEMA 17 format stepper motor (StepperOnline) designed specifically for 1/16” ID 1/8” OD fluidic tubing. Its rotor was 3D printed via selective laser melting of 316 stainless steel (PCBWay). The pump rollers were 3D printed via selective laser sintering from nylon and coupled using ball bearings. A top bracket for roller alignment was laser cut from 1008 steel (SendCutSend). An alignment bracket to hold the tubing and align the pump backing was CNC milled from C360 brass (PCBWay) and coupled to a pump backing. The pump tubing was 1/16” ID” inch OD Pharmed BPT tubing with stops made from 1/8” ID polyvinyl chloride tubing affixed using Loctite 7701 and 4011. Bill of materials in Table S4.

### Fluidic control and automation

Peristaltic pumps were driven by a Pololu Tic T825 stepper motor controller operating at 24 V (Fig. 3b(iv)) controlled over USB using the Pololu ticcmd command line utility (Fig. S13). A common 24 V power supply was used to drive all pumps. Bill of materials in Table S5. Media change, pump priming, perfusion and fluidic scheduling operations were automated using Bash shell scripts running on a PC (Fig. 3b(ii)). Manual override of fluidic operations was implemented using scripts launched with an Elgato Streamdeck (Fig. 3b(i)) via the Core447 StreamController utility.

### Temperature regulation

The apparatus used custom CNC milled 6062 aluminium heat blocks for temperature regulation of the well (Fig. 3b(xi), Fig. S14) and gas exchanger (Fig. 3b(ix)). Omega CN401-11445 PID process controllers (Fig. 3b(iii)) were used for temperature feedback control due to their reliability and safety features, particularly their firmware loop break detection. The controllers received input from type K thermocouples. They controlled heater power via solid state relays (Omron) (Fig. 3b(vii), Fig. S15). The heating system was protected against thermal runaway primarily by the PID controller’s safety routines and secondarily by a thermal fuse mounted in a selective laser melting printed aluminium holder attached to the heat plate assembly with thermal adhesive (MG Chemicals 8329TCM). Commercial 93*×*10 mm resistive heaters operating at 24 V (Generic, e.g. Amazon B09RWXXVN7) served as heating elements. Bill of materials in Table S6.

### Circulatory loop assembly

To create our one-circuit incubator-free recirculatory apparatus, the loop was assembled as in Fig 3a, with further detail in Fig. S16 and Table S7. A swabbable luer lock quick disconnect serving as an access port (Fig. 3a(i)) for the media supply syringe (Fig. 3a(ii)) was coupled to a length of KFlex tubing by means of a luer to 1/16” hose barb connector. This tubing was threaded through a hole drilled in the side of a Peltier device-based desktop refrigerator. Using a 1/16” hose barb elbow fitting, this media supply line was coupled to the Pharmed BPT tubing forming the actuator tubing of a first *media change peristaltic pump* (Fig. 3A(iii)). The outflow was coupled to a length of KFlex tubing containing an interposed check valve (Fig. 3a(iv)) oriented as to only allow fluid transit under outgoing flow from the media supply syringe. This check valve was coupled via a tee junction to the *media recirculation circuit*. This circuit consists of lengths of KFlex tubing. The *afferent leg* (Fig. 3a(v)) of this circuit passes from the supply tee junction and joins the inflow port of the media side of the gas exchanger via a coupler assembled from mating luer lock barb adapters. The circuit, with its contained media having passed through the gas exchanger media side, continues via another length of KFlex tubing coupled proximally to the gas exchanger via another pair of mating luer lock barb adapters and distally by a male luer lock barb adapter with a free spinning collar. This completes the *afferent leg* of the recirculation circuit. This is the point at which the well will be interposed, its inflow port mating to this adapter (Fig. 3a(vii)). Absent the well, the male luer adapter on the distal end of the *afferent leg* is coupled to another identical male luer lock barb adapter via a female-to-female luer lock adapter. With the well present, the two male luer lock barb adapters will be connected to the inflow and outflow ports of the well. From the second male luer adapter continues the *efferent leg* of the recirculatory loop (Fig. 3a(viii)). This leg links the outflow port of the well to the inflow of a second peristaltic pump, the *recirculation pump* (Fig. 3a(xi)). Interposed in this leg is a tee junction to allow ejection of distal-to-well media effluent (Fig. 3a(x)) via the efferent leg of the circuit by volume displacement when fresh media is injected. An outflow check valve (Fig. 3a(ix)) protects this junction from backflow. Finally, the outflow from the peristaltic pump links to the inflow tee junction, completing the circuit.

### Device sterilization

KFlex tubing, gas exchangers and oxygen sensor spots were sterilized in situ by circulation for one hour with 0.2% peracetic acid solution prepared fresh by dilution of 32% peracetic acid (Sigma-Aldrich 269336) in sterile water. All other parts were sterilized by autoclave at 121 °C, 15 psi, 30 min on a vacuum cycle prior to assembly or use.

### Assembled apparatus preparation for experiments

If oxygen or pH sensor calibration was performed, these were done prior to sterilization. After in situ sterilization, the acid was neutralized and the loop was rinsed with a minimum of five loop volumes of sterile PBS. Immediately prior to starting culture, the loop was primed with fresh cell culture media. Access ports were sanitized prior to each access by physical wiping with Oxivir or isopropanol wipes and spraying with 70% isopropanol.

### Gas control

To achieve media dissolved gas control via diffusion across the PMP membrane of the gas exchanger device, an aqueous gas control buffer was prepared. First, in order to achieve oxygen equilibration with room air at cell culture temperature, deionized water was heated for a minimum of one hour at 37 °C in an open vessel under vigorous magnetic stirring. To this was added 14.8 mg/mL sodium carbonate and 67.2 mg/mL sodium bicarbonate and the mixture was stirred in a closed container with minimal head space until fully dissolved. With this freshly prepared mixture, the gas control buffer compartment (Fig. 3b(iv)) was manually syringe filled through a remote fill port attached to the gas exchanger using KFlex gas impermeable tubing. The fill port was sealed using a swabbable self-sealing female luer port (Qosina). The opposite port of the gas exchanger was open to air via an overflow drain tube constructed from KFlex tubing. This was cross-clamped after filling. Replacement of the gas control buffer was achieved by withdrawal through the same remote fill port.

### Loading of organoids

Organoid wells with preinstalled gaskets (Fig. 3g-k)) were autoclaved and placed in clamping elements. Organoids were transferred into the growth chamber well by pipetting with a wide bore pipette tip in a biosafety cabinet. The organoid well was partly filled with media, then closed with screws loosely holding the lid in place. The well assembly was removed from the biosafety cabinet and an automated torque screwdriver (Ingersoll Rand QXFD series) set to 2 N-m was used to tighten the clamping screws to seal the well gaskets against the clamping element as shown in Fig. 3g-j. The procedure is detailed in Fig. S17. The sealed well was placed in the heat block as shown in Fig 3k. At this point, the interior of the well assembly remains aseptic while the exterior no longer requires aseptic handling. Description of required equipment in Table S8.

The luer quick disconnects on the top of the well (Fig. 3j) were sanitized by swabbing with isopropanol wipes. The inflow and outflow lines of the circulatory loop, which had been joined by a sterile coupler and cross-clamped, were removed from the coupler, sprayed with 70% isopropanol and installed into the luer quick disconnects (Fig. 3k). The inflow and outflow lines used male luer lock to barb adapters with free-rotating lock collars to prevent twisting of the lines on installation. After connection, the cross-clamp was removed to allow circulation through the loop, and the residual air volume in the well caused by its partial filling was cleared using the priming subroutine (Fig. 3f) of the fluidic controller.

### pH measurement

For pH measurement free of sensor drift or electrical interference, pH was measured by continuous absorbance spectroscopy using an ams-OSRAM AS7262 spectral sensor on an I^2^C breakout board (Adafruit, 3779) with data recording via a Python script reading the I^2^C data through an Arduino Uno R3. A custom flow cell cuvette CNC milled from polycarbonate, masked with a light-reflecting inner layer of white electrical tape and an outer layer of black electrical tape (3M), illuminated with the white LED onboard the Adafruit breakout and placed inline in the perfusion loop immediately after the gas exchanger. Bill of materials in Table S9.

pH was calculated from measurements using a calibration curve fit by linear regression on the ratio of absorbance on the 570 nm channel to absorbance on the 450 nm channel of the AS7262 sensor. Commercial phosphate buffer standards at pH 7.0, 7.4, 8.0 (Fisher Sci SB108500, SB110500, SB112500) with added 8.0 mg/mL phenol red (to match post-mixing dilution of standard 8.1 mg/L phenol red in media) were used for calibration.

### O_2_ measurement

Oxygen was monitored using a custom polycarbonate flow cell containing an oxygen-sensitive dye spot operating on the principle of oxygen quenching phase fluorometry. We used a chemical resistant oxygen spot purchased from Spectrecology and a Fluorometrics DOPO-2 fiber optic oxygen sensor for phase fluorometry. Data were recorded using the accompanying proprietary software. To avoid motion artefact, the fiber optic cable was rigidly mounted to the cuvette using a custom steel fixturing plates. The thermistor for temperature compensation was affixed to the outside of the fluidic tubing to avoid introduction of a foreign body into the fluidic circuit; this likely results in increased thermal artifact due to the independent effect of temperature of oxygen sensitive dye fluorescence lifetime measurements.

The oxygen sensing flow cell was calibrated by perfusing with freshly prepared aqueous 1% sodium sulfite with catalytic cobalt(II) chloride as a 0% oxygen standard and water equilibrated with room air at 37°C overnight under vigorous magnetic stirring as a 20.9% oxygen standard.

### Evaporation and salt concentration measurement

Media sodium concentrations were measured using a Beckman Coulter Vi-Cell MetaFLEX bioanalyte analyzer. Hypernatremia served as a quantitative proxy for water loss via evaporation, as Weigmann [72] demonstrated that sodium ion concentration in evaporation-susceptible environments accurately reflects evaporation in cell cultures.

Since dissolved electrolytes should not enter gas phase at physiological temperature, sodium concentration change was assumed to reflect estimate global relative salt concentration change. Volume change was calculated as the volume of free water required to produce the observed sodium concentration assuming fixed sodium content. Assuming constant temperature, humidity, advection rate and interfacial surface area, the evaporation rate will be constant, causing salt concentration to vary as <u>^1^</u> and volume to vary as 1 *−Bt*. Data were analyzed by fitting curves using least squares regression on these reciprocal linear and linear relationships, respectively. Data were compared by *t* -tests on curve coefficients *A*, *B*.

### Human induced pluripotent stem cell culture

Human induced pluripotent stem cells (hiPSCs), line WTC11, bearing a constitutively expressed tdTomato fluorophore, were a gift from the Nowakowski lab (University of California, San Francisco). hiPSCs were cultured in a Matrigel (Corning, 354277) coated plate with mTESR1 Plus media (Stemcell Technologies, 100-0276), media changed daily and passaged with ReLeSR (Stemcell Technologies, 100-0484).

### Human vascular organoid generation

Human vascular organoids were generated from hiPSCs following Wimmer et al.’s protocol [58] (Sigma-Aldrich, A6964). Briefly, hiPSCs were detached with Accutase, 12,000 cells were seeded per well of an ultra-low attachment 96-well plate with 10 µM of ROCK inhibitor Y-27632 (10 µM) (Tocris Bioscience, 1254). After 24 hours, a complete media change was performed with mesoderm induction media consisting of N2B27 medium supplemented with CHIR99021 (12 µM) (Tocris Bioscience, 4423) and BMP-4 (2 µM) (Miltenyi Biotech, 130-111-164) for 3 days. Subsequently, media was changed to vascular induction media consisting of N2B27 media supplemented with VEGF-A (100ng/mL) (Peprotech, 100-20) and forskolin (2 µM) (Sigma-Aldrich, F3917), and maintained for another 3 days until vascular sprouts were observed [58]. These resulting vascular organoids were individually embedded in 30 µl droplets of Matrigel on ice and then incubated at 37°C for 1 hour to allow solidification of the Matrigel around the vascular organoid. Embedded organoids were transferred to ultralow attachment 6-well plates and cultured in *VO maintenance media* consisting of StemPro-34 media (Gibco, 10640) supplemented with VEGF-A (100 ng/mL), FGF-2 (100 ng/mL) (Stemcell Technologies, 78003) and 15% fetal bovine serum (Gibco, A525680) and maintained on an orbital shaker at 90 rpm until further use, with media changes every 2 days.

### Human vascular organoid culture in our system

Day 45 human vascular organoids were embedded in 5% gelatin-methacrylate (GelMA) (Cellink, IKG125000005) mixture with 0.5% Lithium phenyl-2,4,6-trimethylbenzoylphosphinate (Cellink, 5269) photoinitiator for crosslinking. In each well of a two-well version of our device, 100 µl of GelMA mixture was added and overlaid with a 45 day old vascular organoid and overlaid on top with 100 µl of GelMA mixture and UV crosslinked for 30 sec. 200 µl of *VO maintenance media* was added and the device was sealed and loaded into our system as described previously. Automated media changes were every 2.5 days using *VO maintenance media*. The vascular organoid was cultured for 3 weeks with hourly tdTomato epifluorescence imaging at 37 Z-stack levels separated by 50 µm as described below.

### Live cell fluorescence imaging of human vascular organoids

A miniature automated fluorescence microscope (YR and DE, in preparation) was built, using an Allied Vision Alvium USB CMOS camera, 10*×* microscopy objective and bespoke lens barrel adapter and vertical positioner (Fig. 4b). Samples were illuminated with a Thorlabs MINTF4 fiber optic light source, Chroma ET560/40x excitation filter, 600nm longpass dichroic mirror (Edmund 69-868) and Chroma ET630/75m emission filter. This was used for time series epifluorescence microscopy of the vascular organoid with 37 Z-stack levels at 1 hour intervals through the well bottom. Z-stack images were aligned within timepoints and brightnesses and gamma parameters were scaled uniformly between all images all time points using batch processing with a custom python pipeline built around the NumPy and Pillow Python libraries. Lone bright pixels attributable to systematic differences in pixel response over long exposures were removed. At each time point, stacks were compared pixel by pixel using NumPy vector operations in order to identify pixels consistently displaying brightnesses inconsistent with their nearest neighbors across the majority of images. These bright “stuck pixels” were removed by replacement with the average of their immediate neighbors as to not produce artefactual images on Z-stack layer alignment or EDF fusion. Next, images were then aligned within Z-stacks using OpenCV’s Enhanced Correlation Coefficient algorithm to correct for any alignment artefacts from Z-stacking.

Extended depth of field (EDF) fusions of the Z-stacks were generated using a Python implementation of complex wavelet transform [73] (Fig. 4c). Their approach was modified to fuse multiple highest scoring pixels as this produced smoother results on blood vessels crossing Z-planes. TIFF stacks were loaded and converted to NumPy arrays, then decomposed into wavelet coefficients using PyWavelets. Db3 Daubechies wavelets with 11 decomposition steps were used. For each pixel, the magnitude of the wavelet coefficients were compared across all images in the stack, and the pixels with the top N coefficients were averaged. These operations were performed using NumPy vector operations where possible.

To obtain stack average datasets for fluorescence signal summation, Z-stack photosets were loaded with Pillow and averaged pixelwise using NumPy vectorized operations. Precise acquisition timepoints were extracted from microscopy image metadata. Growth curves (Fig. 4d) were calculated by summation across pixel-wise averaged Z-stacks. Fluorescence signal was summed across pixels using NumPy. Plots were created using matplotlib, using scipy for statistical analysis.

### Murine embryonic stem cell culture

Cortical organoids were derived from BRUCE4 C57BL/6 murine embryonic stem cells (mESCs). mESCs were expanded and maintained on tissue culture plates coated with 0.5 µg/mL vitronectin (10 minutes in PBS at 37 °C). *mESC maintenance medium* was composed of Glasgow Minimum Essential Medium (GMEM, Gibco 11710) supplemented with 10% ES cell-qualified fetal bovine serum (Gibco 10439), 0.1 mM MEM non-essential amino acids (Gibco 11140), 1 mM sodium pyruvate, 2 mM GlutaMAX (Gibco), 0.1 mM *β*-mercaptoethanol and 0.05 mg/mL Primocin (InvivoGen). Medium was supplemented with 1000 U/mL ESGRO recombinant mouse leukemia inhibitory factor (LIF, Chemicon ESG1107) at each media change. Medium was changed daily. Cells were passaged using TrypLE Express without phenol red (Gibco, 12604) following the manufacturer’s protocol. Cryopreservation was in Bambanker cell freezing medium (Bulldog Bio) in liquid nitrogen vapor phase.

### Murine cortical organoid generation

Dorsal forebrain cortical organoids were generated from mESCs following a modified Hernandez et al. [59] protocol. mESCs, dissociated in 750 µL TrypLE Express were aggregated in PrimeSurface 96 Slit-well plates (S-Bio MS9096SZ) at a density of 3000 to 3500 cells in 100 µL media per well. Aggregates were maintained in 15 mL *mESC maintenance medium* per plate with 10 µM Rho Kinase Inhibitor Y-27632 (Tocris 1254) and 1000 U/mL LIF. At 24 hours, media was exchanged to *forebrain patterning medium*: DMEM/F12 with GlutaMAX (Gibco 10565), 0.5*×* MEM Non-Essential Amino Acids, 0.5 mM sodium pyruvate, 0.5*×* Chemically Defined Lipid Concentrate (CD Lipid, Gibco 11905), 2*×* B-27 without Vitamin A (Gibco 12587), 0.1 mM *β*-mercaptoethanol and 0.05 mg/mL Primocin. Media was changed daily. To promote dorsal forebrain patterning, 10 µM Y-27632, 5 µM XAV939 WNT pathway inhibitor (StemCell), 5 µM TGF-*β* receptor inhibitor SB431542 (Tocris 1614), and 250 nM LDN193189 BMP inhibitor (StemCell) were added to the medium.

On day 6, organoids were transferred to ultra-low adhesion 6-well plates (Corning 3471), with 3 mL fresh *progenitor expansion medium* per well. Medium composition was modified from [59]: DMEM/F12 with Glutamax mixed 1:1 with BrainPhys Neuronal Medium (StemCell 05790), 2*×* B-27 without Vitamin A, 1*×* MEM Non-Essential Amino Acids, 1*×* CD Lipid, 0.64 mM GlutaMAX, 0.05 mg/mL Primocin and 200 µM ascorbic acid. From days 6-12 organoids were cultured under orbital shaking at 69 rpm with media changes every 48 hrs.

### Cortical organoid maintenance

From days 13-25, medium was *neuronal maturation medium* consisting of BrainPhys Neuronal Medium, 2X B-27, 1*×* CD Lipid, 5 µg/mL porcine heparin (Sigma-Aldrich H3149) and 0.05 mg/mL Primocin. 200 µM ascorbic acid was added until day 25.

Media changes were 2 days. On Day 26 2*×* B-27 was replaced with 1*×* B-27 Plus supplement (Gibco A35828).

### Culture of cortical organoids in our system

Organoids were divided into three equal cohorts for trials comparing incubator culture in static well plates, incubator culture in well plates under inertial agitation on an orbital shaker and culture in our incubator-free recirulatory system at a flow rate of 1 mL/min. Trials lasted 5.5 days, with media changes on days 2 and 4. Media changes were performed manually in shaker in incubator conditions and by fluidic automation in our recirculatory apparatus. On Day 5.5, organoids were recovered from each condition and subcohorts from each subjected to the following analyses:

### Metabolic activity quantification of cortical organoids

Cell viability was assessed using an XTT (2,3-Bis-(2-methoxy-4-nitro-5-sulfophenyl)-2H-tetrazolium-5-carboxanilide) colorimetric assay, which measures total organoid mitochondrial NAD(P)H dependent oxidoreductase activity as a proxy for cell proliferation and viability. From each experimental condition, single organoids were transferred to wells of 24-well plates. Organoids were washed with 1*×* PBS. 700 µL of phenol red free DMEM without glutamine (Gibco 31053) supplemented with GlutaMAX and 0.05 mg/mL Primocin was added to each well. Next, 300 µL of freshly mixed working solution containing 7.68 µg/mL phenazine methosulfate (PMS) and 25.2 mg/mL XTT in the same media was added to each well. Plates were incubated for 4 hours 45 minutes at 37 °C, 5% CO2. XTT formazan conversion was measured via absorbance at 475 nm using an Ocean Insight FLAME-T-Vis-NIR-ES spectrophotometer in fresh disposable polystyrene cuvettes in an Ocean Optics Square One cuvette holder with background correction via subtraction by measured absorbance at 660 nm. We opted for this measurement setup over a plate reader assay due to superior technical replicability.

We used a linear mixed effects model for comparisons treating experimental cohorts as random effects. Outliers measuring twice their condition median were excluded as attributable to fusion of two organoids. Data were evaluated by pairwise *t*-tests for noninferiority of our apparatus using a 10% relative noninferiority margin, justified by high inter-cohort variability in organoid culture as demonstrated in (Fig. 5c).

### Immunohistochemistry of cortical organoids

Whole-mount analysis was followed the protocol described in [59]. Organoids were fixed in 4% paraformaldehyde at 4 °C overnight, embedded in 4% low-melt agarose (Invitrogen 16520-050), and sectioned to 40 µm thickness using a Leica VT1000S vibratome. Sections were incubated for 1 hour at 4 °C in a blocking buffer containing 5% donkey serum, 1% BSA, and 0.5% Triton X-100. Primary antibodies were then applied in a blocking solution of 2% donkey serum, 0.1% Triton X-100 and incubated overnight at 4 °C. Primary antibodies used were: rabbit anti-Map2 (Thermo Fisher 17490-1-AP), mouse anti-Sox2 (Santa Cruz Biotech sc365823), mouse anti-Tubb3, 1:2000 (Biolegend 801201), rabbit anti-POU3F2/Brn2, 1:400 (Invitrogen PA530124), mouse anti-Gfap, 1:400 (Sigma-Aldrich G6171), rabbit anti-GABA, 1:375 (Thermo Fisher PA5-32241). Primary antibodies were detected with donkey anti-rabbit Alexa Fluor 546, 1:750 (Invitrogen A10040) and donkey anti-mouse Alexa Fluor 647, 1:750 (Invitrogen A31571) applied overnight at 4 °C and counterstained with 3.5 µM DAPI (Invitrogen D1306). Samples were mounted using Fluoromount-G (SouthernBiotech 010001) prior to imaging. Marker selection was guided by prior characterizations of mouse dorsal cortical organoids [59].

Organoid imaging was performed on a Zeiss AxioImager Z2 microscope using 10*×* and 20*×* objectives with ZEN Blue software (Zeiss). For each organoid imaged at 10*×* objective, z-stacks were acquired at 1 µm intervals across 37 optical planes, and each plane was tiled into four fields of view and stitched in the ZEN software. From these stacks, 13 representative planes were selected and merged into a maximum-intensity z-projection composite using ImageJ. Projected images were split into individual fluorescence channels, adjusted for brightness and contrast (equally between experimental and control conditions without nonlinear adjustments), converted to RGB format, saved as TIFF files, and recombined into composite figures. For 20*×* acquisitions, z-stacks spanning 20 µm were collected at 2 µm intervals (11 planes total). For figure presentation, the most in-focus optical plane was selected, while the z-stack was used to navigate through the tissue and examine specific regions of interest. All images were cropped to the same field of view to ensure consistent scaling across conditions. Scale bar dimensions were calculated in ZEN Blue software using microscope metadata.

### Live cell brightfield imaging of cortical organoids

A miniature custom automated brightfield microscope was built, using an Allied Vision Alvium USB CMOS camera, a 4*×* microscopy objective and bespoke lens barrel adapter and vertical positioner. This was based on technology described in [74]. This was used for time series light microscopy of organoids at 2-hour intervals with 21 Z stacks separated by 10 µm intervals through the well bottom. Images were aligned, background masked (to remove well boundary), cropped and brightness normalized uniformly in batch using a custom python pipeline built around the NumPy and Pillow libraries. EDF fusions were generated using a custom Python implementation of complex wavelet transform [73] method with Daubechies db3 wavelets with fusion of the top two ranks. Growth curves were calculated on pixel-wise averaged Z-stacks using the cropped and masked data to avoid spurious counting of background shadowing.

### Electrophysiology of cortical organoids

Day 33 mouse cortical organoids, grown for 5.5 days either with our system or on shaker in the incubator, were plated on MaxOne high-density multielectrode arrays (HD-MEAs; Maxwell Biosystems) following the protocol in [59] (Fig. 6a). HD-MEAs were prepared by coating with 500 µL of 80 ug/mL laminin in PBS overnight at 37 °C. Organoids were cultured for 8 days on the HD-MEAs to allow attachment and adaptation to the plate surface. Electrophysiological recordings were acquired on organoid day 41.

Initial plate scanning was performed in sets of 1024 electrodes arranged in a rotating checkerboard pattern across the 26,400-electrode array, with each configuration recorded for 30 seconds. The 1020 most active electrodes from this scan forming a subset pairwise separated by a minimum of 50 µm were selected as putative neuronal units for further analysis. Recordings from this subset were performed in *neuronal maturation medium* in a humidified incubator maintained at 37 °C with 5% CO_2_, using a 20 kHz sampling rate. Raw data were bandpass filtered to 300–6000 Hz and processed for spike sorting with Kilosort2 [75] via a custom Python-based pipeline [69]. Putative neuronal units were excluded from analysis if they exhibited interspike interval violation rates *>* 0.5, firing rates *<* 0.1 Hz, or signal-to-noise ratios *<* 3 [76].

Spatial distribution of neuronal units was visualized using bubble maps with bubble size proportional to the peak-to-peak amplitude of each unit’s extracellular waveform (capped at 50 µV) Units were color-coded by cluster identity. Black bounding boxes indicate selected regions of interest for detailed waveform analysis. Bubble sizes are proportional to the amplitude of observed extracellular action potentials (maximum bubble size truncated to 50 µV amplitude). Within selected bounding boxes, spatial footprints of neuronal units were displayed by overlaying their measured waveforms according to electrode position. Each unit’s footprint included waveforms from both center and neighboring electrodes, color-matched to the amplitude map. Center electrodes were indicated with black-edge markers.

### Hierarchical clustering of STTC-calculated connectivity

Network synchrony was evaluated using spike-time tiling coefficient (STTC) matrices calculated with the Catalogger framework [70]. For each recording, the 60 most active units were selected based on firing rate, and pairwise STTC values were computed with a temporal window of Δ*t* = 10 s. STTC similarity matrices were converted to dissimilarities by computing 1 *−* STTC, symmetrized, and zeroed along the diagonal. The resulting pairwise distances were subjected to agglomerative hierarchical clustering (average linkage; scipy.cluster.hierarchy). Dendrogram leaves were used to reorder the STTC matrix for visualization as a clustered heatmap, grouping units with similar temporal firing relationships. This method was conducted in line with previously published applications of hierarchical clustering on STTC matrices for functional connectivity analysis in human hippocampal slices [77].

### Waveform clustering and ISI histogram analysis

Average spike waveforms for each unit were extracted from the qm.npz files and flattened for dimensionality reduction. To visualize waveform diversity and group similarity across culture conditions, waveforms were standardized using StandardScaler and projected into two-dimensional space using Uniform Manifold Approximation and Projection (UMAP; n neighbors = 15, min dist = 0.1, metric = “euclidean”). K-means clustering (n = 7) was applied to the UMAP embedding to assign waveform clusters. Units were labeled by culture condition based on metadata and visualized as triangles (our system) or circles (shaker in incubator). Cluster colors were assigned using a custom categorical palette (CUSTOM PAL SORT 3) with consistent ordering across conditions. Code for UMAP projection, clustering, and visualization was adapted from previously published multimodal electrophysiology pipelines by Andrews et al. [78] and Geng et al. [69].

For each UMAP-derived cluster, we plotted the average extracellular waveform template by computing the mean across all units assigned to that cluster. Waveform traces were normalized by their local maximum to facilitate comparison across clusters. To examine temporal firing patterns, inter-spike interval (ISI) distributions were calculated for each unit by differencing consecutive spike times. Units with fewer than two detected spikes or producing invalid ISI values were excluded from analysis. ISI values were binned from 0 to 100 ms using 30 uniform bins, which provided sufficient resolution to distinguish bursting activity (short ISIs) from tonic firing (longer ISIs). Histograms were generated separately for organoids matured in the incubator-free perfusion system and the shaker in incubator condition, enabling comparison of firing statistics across conditions. Cluster-level ISI histograms thus represent the pooled distribution of spike intervals from all units within the same cluster and culture condition.

