## Supplemental Materials for "Incubator-Free Organoid Culture in a Sealed Recirculatory System"

### Supplemental Material for *Incubator-Free Organoid Culture in a Sealed Recirculatory System*

#### Supplemental Results

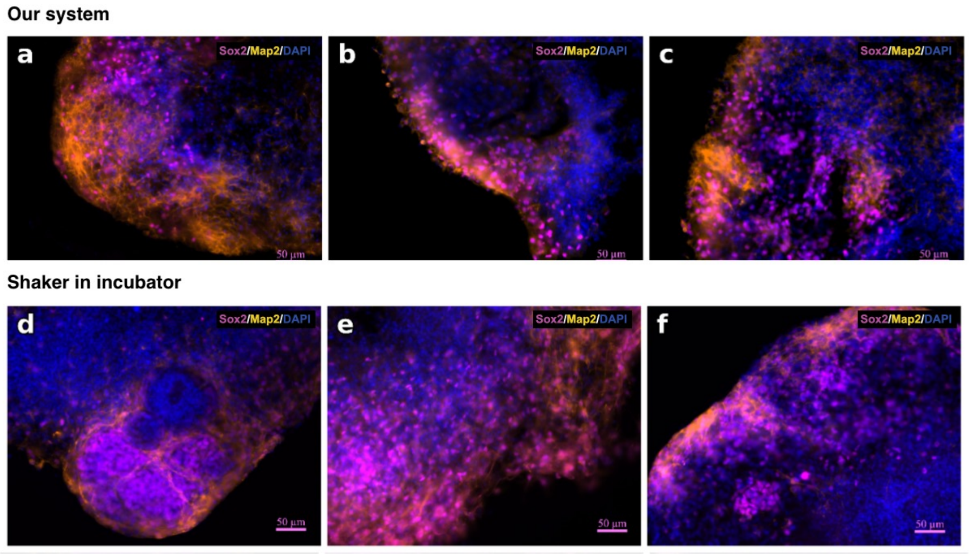


**Fig. S1 Preserved dorsal forebrain organization demonstrated in additional mouse organoids cultured in our system and in standard shaker in incubator conditions** Organoids were stained for Sox2 (magenta, neural progenitor marker), Map2 (orange, neuronal marker), and DAPI (blue, nuclei). Scale bars 50 µm. Images are representative regions from separate organoids. **a**–**c** Immunofluorescence images of three mouse dorsal cerebral cortex organoids, Day 31-33, cultured for 5.5 days in our system. **d**–**f** Immunofluorescence images of three independent mouse dorsal cerebral cortex organoids, Day 31-33, cultured for 5.5 days on shaker in the incubator.

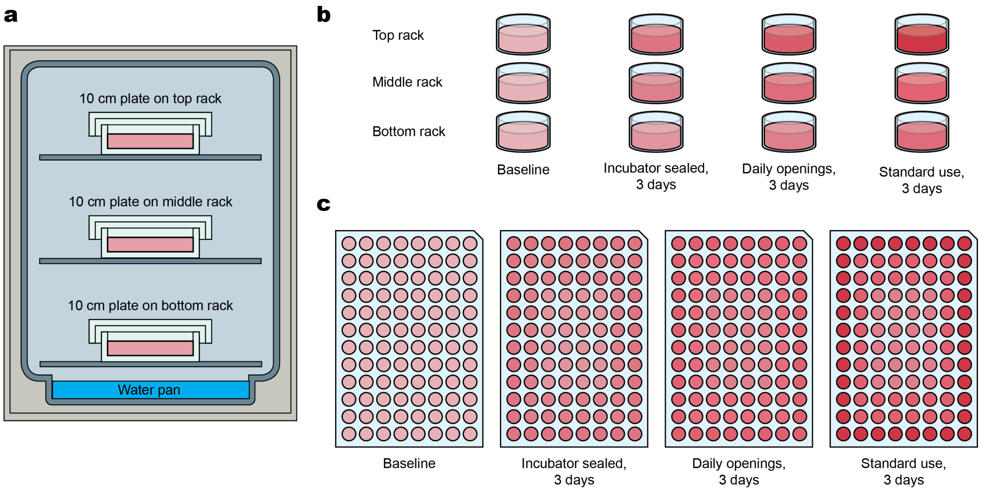


**Fig. S2 Theoretical Schematic of Differential Evaporation Patterns in Incubator Culture.** This figure presents a conceptual overview of differential evaporation severity based on within-incubator and within-plate conditions, based on data collected from our experimental measurements. **a**, Placement of plate within the vertically stacked shelves of a standard incubator affects distance of the plate from the humidifying water bath at the bottom of the incubator and thereby could affect the ambient humidity to which the plate is exposed absent perfect mixing of air. **b**, Cartoon of differences in evaporation corresponding to difference in distance of plate from water bath. Color intensity across top, middle, and bottom incubator racks reflects approximate evaporation rates in 10 cm plates after 3 days under varying incubator usage: sealed, once-daily openings, and standard use (openings as needed for shared cell culture usage in an academic laboratory.) Darker shades indicate higher evaporation, with the top rack consistently showing the greatest loss. **c**, Evaporation distribution across a 96-well plate under the same incubator conditions. Peripheral (edge) wells exhibit substantially higher evaporation than center wells, with severity increasing as incubator openings increase.

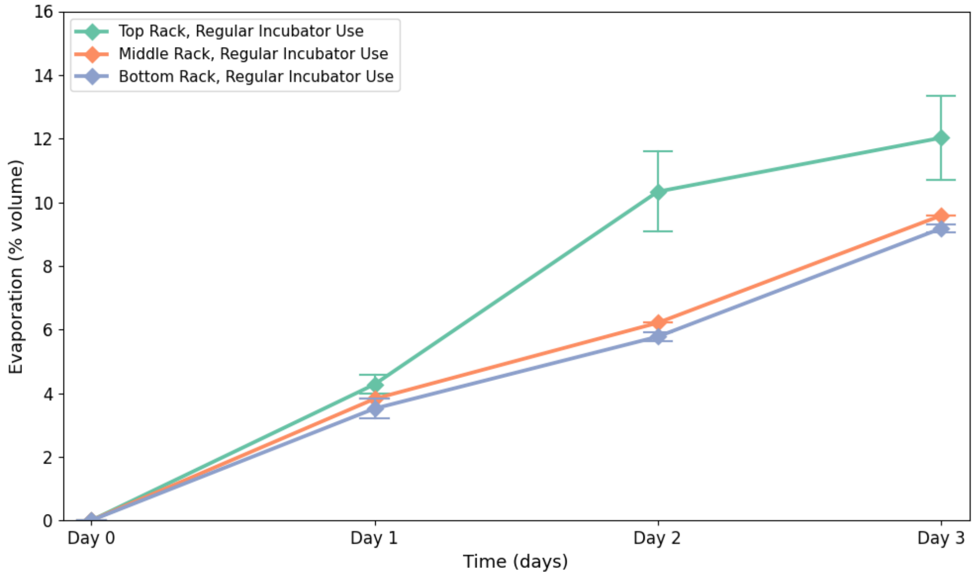


**Fig. S3 Variability of evaporation with respect to plate position for well plates in a humidified cell culture incubator**, assessed by measuring evaporation from plates at different distances from the humidification water bath at the base of the incubator. 10 cm tissue culture plates (Corning 353003) containing 10 mL media were placed on the top, middle and bottom racks of the incubator. At days 1, 2 and 3, plates were sacrificed and their evaporation was quantified by proxy of sodium concentration measured using a Beckman Coulter Vi-Cell MetaFLEX bioanalyte analyzer.

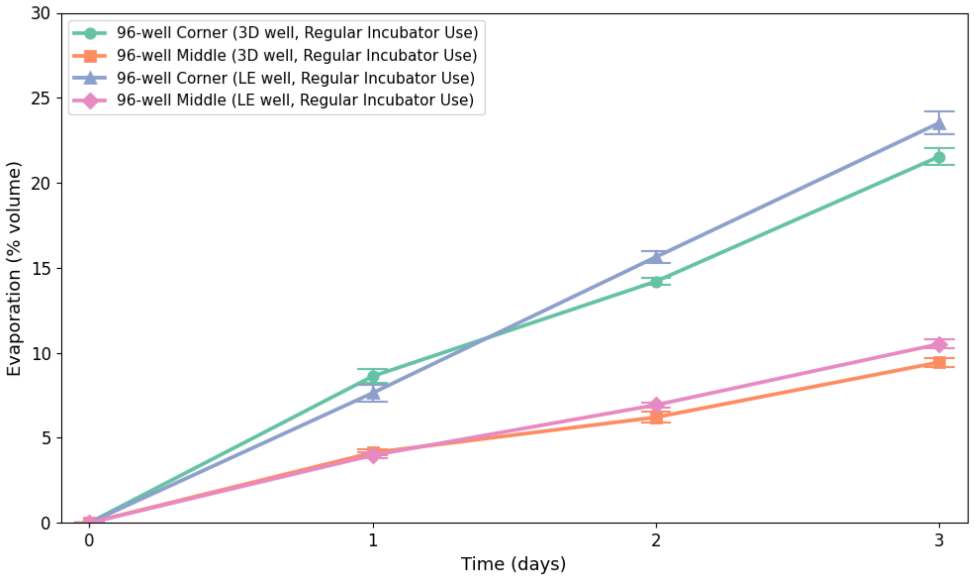


**Fig. S4 Variability of evaporation with respect to well position within well plates and well geometry**, assessed by comparing corner and central wells in two different 96 well plate designs. In both cases, plates were placed on the top rack of the incubator. The two plate designs used are standard within our research group: S-Bio PrimeSurface® 3D Culture Spheroid plates ("3D well", S-Bio MS-9096VZ) and flat bottomed plates with a low evaporation lid ("LE well", Corning 3595). At days 1, 2 and 3, plates were sacrificed and their evaporation was quantified by proxy of sodium concentration measured using a Beckman Coulter Vi-Cell MetaFLEX bioanalyte analyzer.

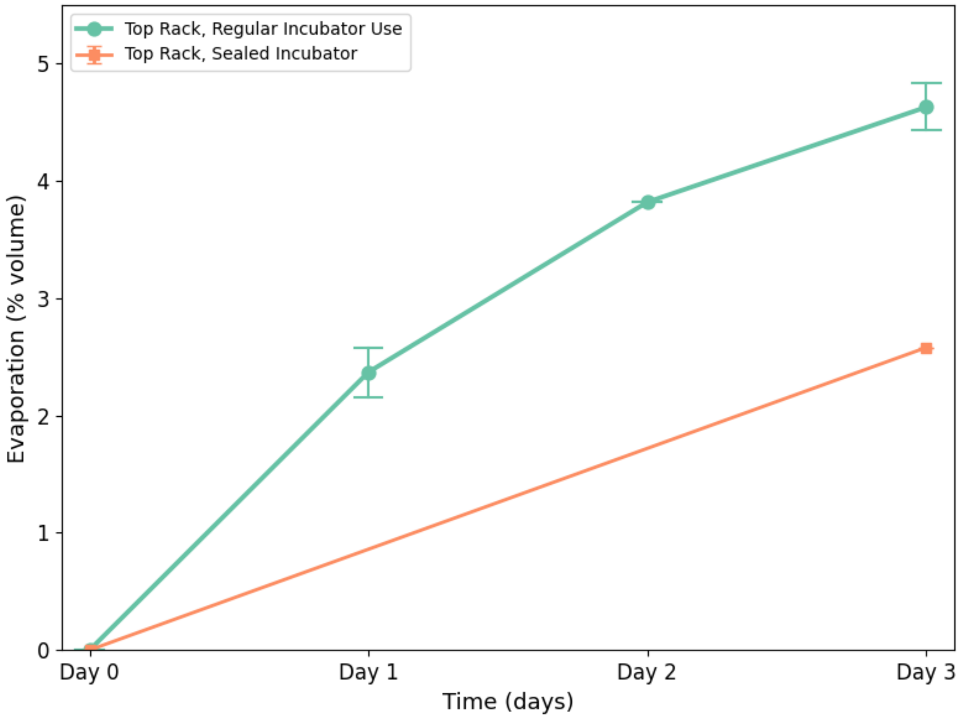


**Fig. S5 Variability of evaporation with respect to incubator access patterns**, assessed by comparing evaporation of media T25 flasks in a sealed incubator which was not accessed for three days versus a shared incubator in the lab routinely accessed throughout the workday. At day 3 for the sealed incubator condition and days 1, 2 and 3 for the regular incubator use condition, plates were sacrificed and their evaporation was quantified by proxy of sodium concentration measured using a Beckman Coulter Vi-Cell MetaFLEX bioanalyte analyzer.

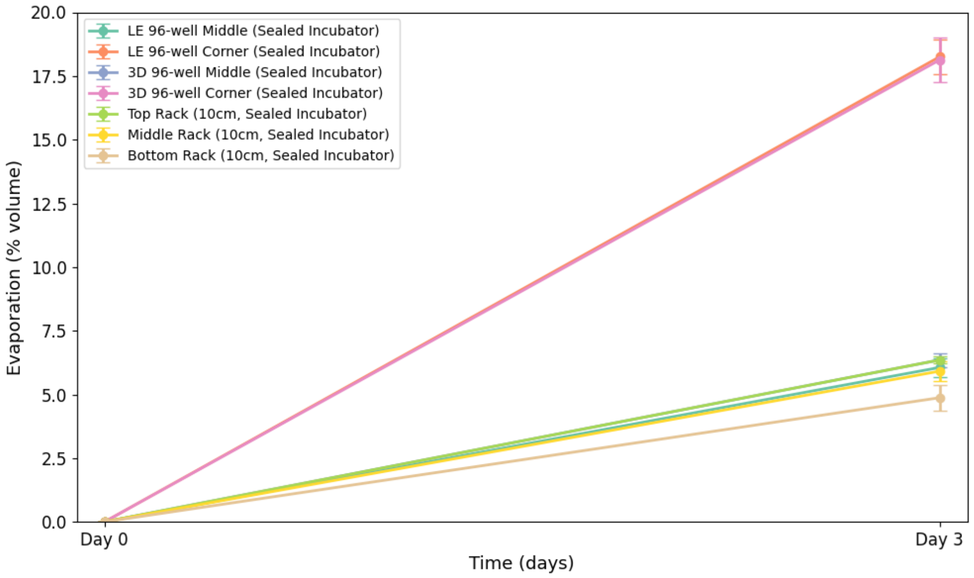


**Fig. S6 Variability of evaporation with respect to plate design in a sealed incubator**, assessed by comparing evaporation of media from 96 well plates ("3D", S-Bio MS-9096VZ and "LE", Corning 3595) as well as 10 cm plates (Corning 353003) in a sealed incubator which was never opened for three days. At day 3 for the plates were sacrificed and their evaporation was quantified by proxy of sodium concentration measured using a Beckman Coulter Vi-Cell MetaFLEX bioanalyte analyzer. Note that evaporation for corner wells of 96 well plates is similar, as well as evaporation for middle wells of 96 well plates and of 10 cm plates on the top rack of the incubator, leading to superimposed data.

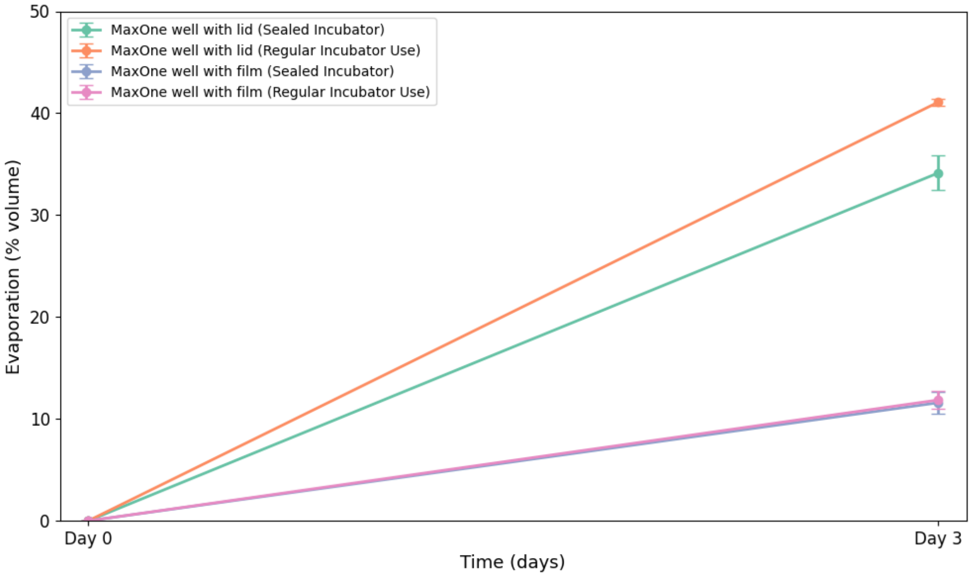


**Fig. S7 Evaporation in Maxwell MaxOne Single-Well High-Density Microelectrode Array electrophysiology plates.** This was performed in both a sealed incubator, never accessed for 3 days and in an incubator under normal use. Lidded wells use the standard lid supplied with the MaxOne wells. Wells with film use 1 mil FEP film (CS Hyde, 23-1FEP-2-50) secured with a 3D printed holder which seals with an elastic silicone O-ring. At day 3 for the plates were sacrificed and their evaporation was quantified by proxy of sodium concentration measured using a Beckman Coulter Vi-Cell MetaFLEX bioanalyte analyzer.

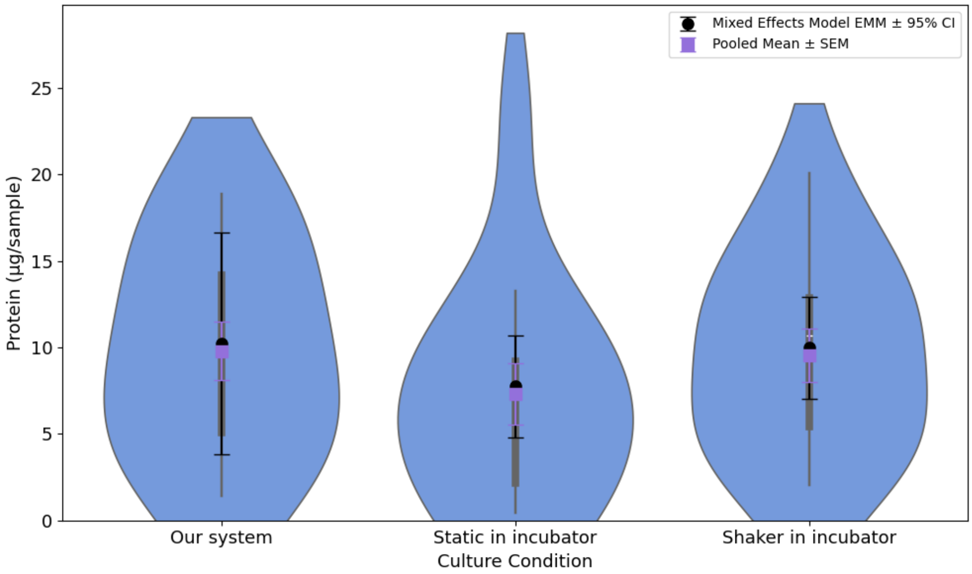


**Fig. S8 Protein quantification of cerebral organoid samples across culture conditions.** Organoids were snap-frozen and lysed using Invitrogen cell extraction buffer (Thermo Fisher, FNN0011) supplemented with 1mM phenylmethylsulfonyl fluoride (Sigma-Aldrich 78830) and 1 mM Sigma protease inhibitor cocktail (Sigma-Aldrich P2714) according to manufacturer instructions, then debris was removed by centrifugation. Total protein was quantified using a detergent-compatible Bradford assay (Thermo Fisher 23246) using absorbance at 595 nm using an Ocean Insight FLAME-T-Vis-NIR-ES spectrophotometer in fresh disposable polystyrene cuvettes in an Ocean Optics Square One cuvette holder. A standard curve spanning 0-1500 µg/mL bovine serum albumin was prepared fresh from kit standard ampules. Data corresponds to the same experimental groups seen in Fig 5c of the main text. Violin plots depict pooled sample distributions, with superimposed boxplots in grey. Pooled means +/- standard errors of the mean (purple) and estimated marginal means ± 95% confidence intervals from a linear mixed-effects model accounting for experimental cohort (black) are overlaid.

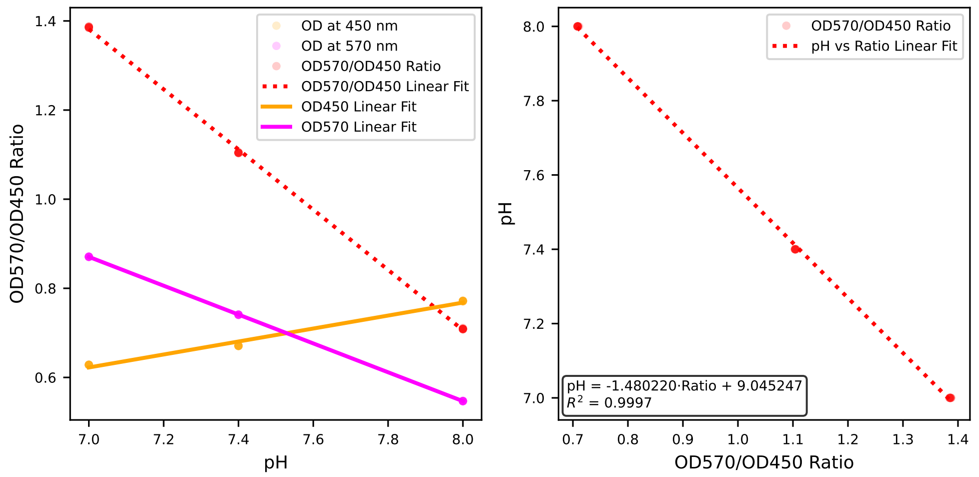


**Fig. S9 Optical pH Standard Curve.** Linear regression on OD570 and OD450 ratio versus pH of phenol red-dyed commercial pH standards yielded the transformation ($pH = -1.480220 Ratio_{OD570/OD450} + 9.045247$) used to generate the pH data shown in the body of this paper. We use this regression model to fit a standard curve within the limited region of pH 7 to 8 spanned by the optical pH standards used. To generate these curves, optical standards prepared from addition of phenol red to pH 7.0, 7.4, and 8.0 phosphate buffer pH standards were prepared to a final target concentration of 7.94 µg/mL, corresponding to 98% of the 8.1 µg/mL concentration present in Gibco DMEM/F12 medium, to account for dilution during media preparation. The sensor was disconnected from the fluidic circuit via its luer fittings without disturbing its position, allowing efficient loading and washout between samples. For each standard, 10 mL of solution was injected through the sensor, and the outflow was collected to ensure complete volume replacement. Tubing was aspirated dry between samples to avoid cross-contamination. Optical density readings were recorded at 570 nm and 450 nm, and a calibration curve was generated by plotting OD570/OD450 ratios against pH, followed by linear regression to determine the line of best fit.

**Table S1** Bill of materials for well manufacturing

| **Component or tool description** | **Specification** | **Product identification and/or source** |
|---|---|---|
| Well body CNC blanks | CNC milled fabrication | 3/8” polycarbonate (McMaster Carr 8574K311), CNC milled on Carbide3D Nomad3 desktop mill |
| Well body CNC fabrication | CNC fabrication, from CNC blanks, using fixturing jig | Milled from well body CNC blanks using Bantam Tools Desktop CNC milling machine and custom fixturing jig |
| Fixturing jig for well body blanks | CNC milled fabrication | 304 stainless steel, PCBWay |
| 3D printer for black resin | Formlabs biomedical 3D printer | Form 3B |
| Black 3D printer resin | Formlabs BioMed Black for Form 3 | Formlabs RS-F2-BMBL-01 |
| 3D printer for optically clear resin | Asiga Ultra 32 | Asiga 07324 |
| 3D printer tray for optically clear prints | Asiga UltraGLOSS build tray | Asiga 05868 |
| 3D printer resin for optically clear prints | Detax Freeprint Ortho 385 | Detax 04095 |
| UV curing chamber for 3D prints | NK Optik Otoflash | NK Optik G171 |
| Fluorosilicone inner gasket | 1/16” 020 size fluorosilicone O-ring | McMaster Carr 8333T13 |
| Perfluoropolymer outer gasket | 1/16” 025 size Aflas O-ring | McMaster Carr 5240T211 |
| Bottom well window | CNC routed polycarbonate | 1/8” polycarbonate (McMaster Carr 8574K26), CNC routed to 20 mm circle |
| Adhesive for window gluing | Loctite 4311 | RS Hughes 079340-00003 |

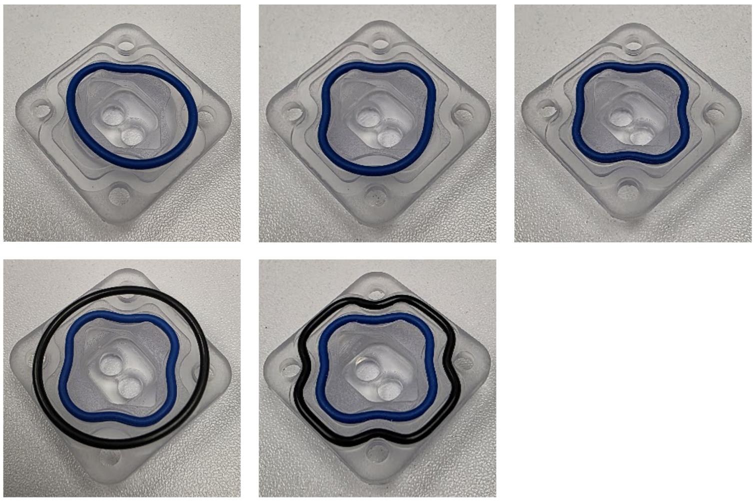

**Fig. S10 Steps in insertion of well gaskets.** Sealing of the wells uses an inner gasket made from fluorosilicone (blue) and an outer gas impermeable Aflas perfluoroelastomer (black) gasket. The cruciform gasket glands are calculated to have the same perimeter as standard-sized O-ring gaskets; the gaskets are fed into the gland to form a cruciform shape, retained by friction against the gland edges. Gasket is progressively fed into cruciform gasket gland. Starting from one corner and working outward, the tension of the gasket against the S-bends will hold the gasket in place.

**Table S2** Bill of materials for well lid manufacturing

| **Component or tool description** | **Specification** | **Product identification and/or source** |
|---|---|---|
| Lid body | CNC milled fabrication | 1/4” polycarbonate (McMaster Carr 8574K281), CNC milled on Carbide3D Nomad3 or Bantam Tools desktop mill |
| Thread mill | 1/4”-28 UNF helical flute thread mill | Micro 100 TM-250-28 |
| Chamfer mill | 90 degree included angle | Harvey tool 47645 |
| Lid white light diffuser | CNC milled fabrication | White 1/8” polycarbonate, TAP Plastics, Tuffak 7328 |
| 1/4”-28 UNF to female luer adapter | Generic, Polycarbonate | Cole Parmer EW-50120-86 |
| Self-sealing disconnect | Silicone swabbable female luer disconnect to male luer lock | Qosina 80213 |
| Adhesive for connectors | Loctite 4311 | RS Hughes 079340-00003 |

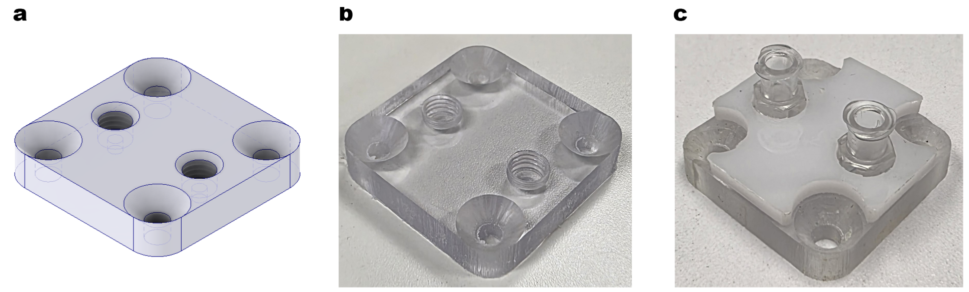

**Fig. S11 Well lid with optional overlaying white light diffuser. a**, 3D CAD file for fabrication. **b**, CNC milled well before insertion of ports or addition of optional white light diffuser. A thread mill is used to thread the ports. c, Addition of luer ports by insertion of $\frac{1}{4}$"-28 UNF to female luer adapters and addition of optional white light diffuser, CNC milled from white polycarbonate.

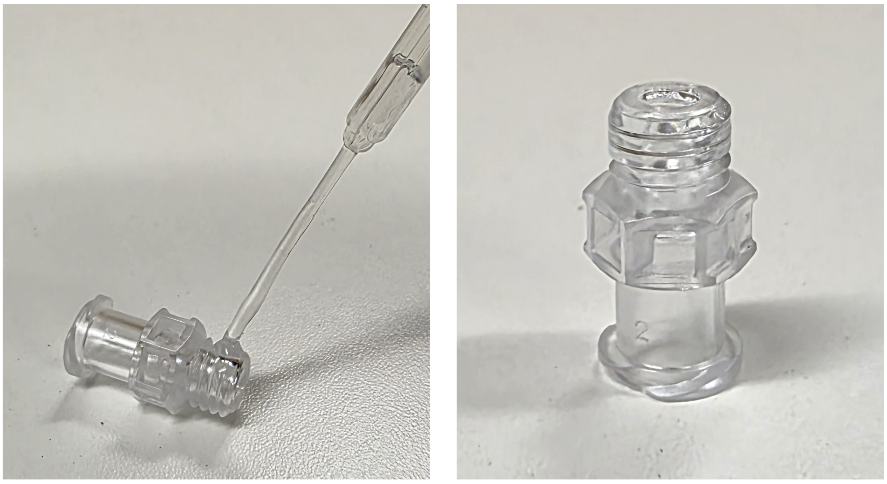

**Fig. S12 Application of Loctite 4311 adhesive to threads of $\frac{1}{4}$”-28 UNF thread to female luer adapters.** This procedure is critical to construction of well lid and gas exchanger reservoir connections. To further prevent bubbling within the thread region, an overnight cure ambient moisture cure is used rather than UV curing. Following this overnight cure, the junction of the adapter with the lid is coated with an overlaying layer of Loctite 4311, taking care to not allow introduction of bubbles (if bubbles form, they can be pierced with a scalpel or deflated with a needle) and then UV cured using a Loctite CL15 UV light source, on rotary turntable. This may be repeated an additional time.

**Table S3** Bill of materials for gas exchanger construction

| **Component or tool description** | **Specification** | **Product identification and/or source** |
|---|---|---|
| Gas exchange media pathway component | CNC milled fabrication | 3/8” polycarbonate (McMaster Carr 8574K311), CNC milled on Carbide3D Nomad3 desktop mill |
| Reservoir component | CNC milled fabrication | 1/2” polycarbonate (McMaster Carr 8574K321), CNC milled on Carbide3D Nomad3 desktop mill |
| Polymethylpentene membrane | Mitsui TPX polymethylpentene, 5 mil | CS Hyde 33-5F-12 |
| Primer for polymethylpentene and tubing | Loctite 7011 | RS Hughes 079340-19886 |
| Adhesive | Loctite 4311 | RS Hughes 079340-00003 |
| Media pathway tubing | 1/16” ID x 1/8” OD x 1/32” Wall PharMed® Tubing | US Plastic Corp 57739 |
| 1/4”-28 UNF to female luer adapter | Generic, Polycarbonate | Cole Parmer EW-50120-86 |
| Reservoir access tubing | Kynar KFlex 1-2, 1/16” ID, 1/8” OD | Eldon James KFLEX1-2 |
| Male luer lock to 1/16” barb adapter | Generic, Polycarbonate | Qosina 11545 or similar |
| Stopcock | F luer to male luer lock | Qosina 99724 |
| Self-sealing disconnect for reservoir access | Silicone swabbable female luer disconnect to male luer lock | Qosina 80213 |
| Deburring tool, square edges | Generic S-blade deburring tool | Amazon B07RM1D6WD or similar |
| Deburring tool, tubing holes | Generic handheld countersink reamer | Amazon B09VS3PS66 or similar |
| Gluing weight | Custom laser cut steel | SendCutSend 0.5” HR A36/1008 mild steel, custom parameters from online part builder: 50mm x 50mm rectangle, 3.9mm rounded corners |
| Rotary turntable | Generic | Amazon B095HKVW46 or similar |

**Table S4** Bill of materials for pump assembly

| Component or tool description | Specification | Product identification and/or source |
|---|---|---|
| Stepper motor | Waterproof NEMA17 | Stepperonline 17IP65-06 or 17IP67-06 |
| Pump body | CNC fabrication | C360 brass, PCBWay, file peristaltic pump v7.ipt |
| Peristaltic pump tubing | 1/16" ID x 1/8" OD x 1/32" Wall PharMed® Tubing | US Plastic Corp 57739 |
| Pump body/frame | CNC fabrication | PCBWay, CNC brass, file peristaltic pump v7.ipt |
| Pump backing | 3D printed | PCBWay, SLS glass fiber nylon |
| Rotor body | CNC fabrication | PCBWay, CNC brass |
| Rotor roller | 3D printed | PCBWay, SLA resin |
| Rotor roller top retaining plate | Sendcutsend, mild steel, brass or stainless | Sendcutsend |
| Rotor body to driveshaft retaining screw | M2 x 4mm grub screw | McMaster Carr 93038A112 |
| Rotor roller retaining screw | M3 x 25 mm button head | McMaster Carr 90910A575 |
| Rotor roller retaining screw threadlocker | Loctite Red | Amazon B000FP8EUS |
| Rotor roller nylon washer | Generic M3 x 6mm x 1mm | Amazon B01N7CJMVL or similar |
| Rotor roller bearing | Generic R63ZZ class 3mm x 6mm x 2.5mm Ball Bearing | Amazon B075CMRGY6 or similar |
| Tubing retainer | Nylon 3D printed fabrication | SLS nylon, PCBWay, file peristaltic pump retainer.ipt |
| Locating pin for tubing retainer | M3x10mm stainless locating pin | Amazon B0CT2DQL31 or similar |
| Locating pin for backing | M3x16mm stainless locating pin | Amazon B0CT2HB3BK or similar |
| Retaining screw for tubing retainer | M4x16mm brass thumbscrew | Amazon B0BRZ3LBYV or similar |
| Compression spring for pump backing | 10 mm long, 0.5mm wire size, 5mm ID compression spring | Amazon B08FDVVTDQ |
| Thumbscrew for pump backing | M4x30mm brass thumbscrew | Amazon B0CR1DT9R9 or similar |
| Body to motor mounting screw | Generic M3 x 25 mm screw | McMaster Carr 92832A234 or similar |
| Tubing retainer - material | Generic 1/8" ID nylon tube | Amazon B07QFMMW3P or similar |
| Tubing retaining tube glue | Loctite 4011 or 4311 | RS Hughes 079340-19886, 079340-18680 |
| Tubing retaining tube glue primer | Loctite 7701 | RS Hughes 079340-00003 |

**Table S5** Bill of materials for pump control

| Component or tool description | Specification | Product identification and/or source |
|---|---|---|
| Motor controller mount | FDM 3D printed, PLA or PCTG | |
| Motor controller | Pololu Tic T825 | Pololu 3130, Digikey 2183-3130-ND |
| Motor controller USB cable | 15-20 cm 90 degree (left angle) micro-USB to USB-A cable | Amazon B0BXY66P5R or similar |
| Motor controller mounting screw | Generic M2 x 8mm cap head or button head screw | McMaster Carr 91290A015 or similar |
| Power supply | Generic 24V 6A or 10A DC power supply | Amazon B0CZ77JSQX or similar |
| Power supply adapter | Generic female 5x5mm x 2.1mm DC jack to screw terminal | Amazon B0C4JJ2HB4 or similar |
| Power splitter for multiple motors | Wago 3- or 5-connector lever splice connector | Wago 221-413 or 221-415, Digikey 2946-221-413-ND, 2946-221-415-ND |
| Inline fuses | Generic 5x20mm format inline fuse holder and fuse, sufficient amperage to allow 2A per motor | Amazon B0813Q4S6P or similar |
| Motor controller internal wiring for motor drive wires | Generic 24 AWG | Amazon B0881H2LR5 or similar |
| Motor drive wire coupler | Wago 221-2401 2-connector lever splice connector | Wago 221-2401, Digikey 2946-221-2401-ND |
| Motor controller internal wiring for power | Generic 20 AWG | Amazon B089CXJPSC or similar |
| Motor controller internal wiring ferrules | Generic 20, 24 AWG | Amazon B0DRJ9CDNG or similar |
| Motor controller wiring stripping tool | With depth gauge | Knipex 121202 or similar |
| Motor controller internal wiring crimping tool | Generic | Amazon B0DJNYNNC2 or similar |
| USB hub | StarTech.com 7-Port USB 2.0 Hub | Startech ST7200USBM, Amazon B003AVPUZG |

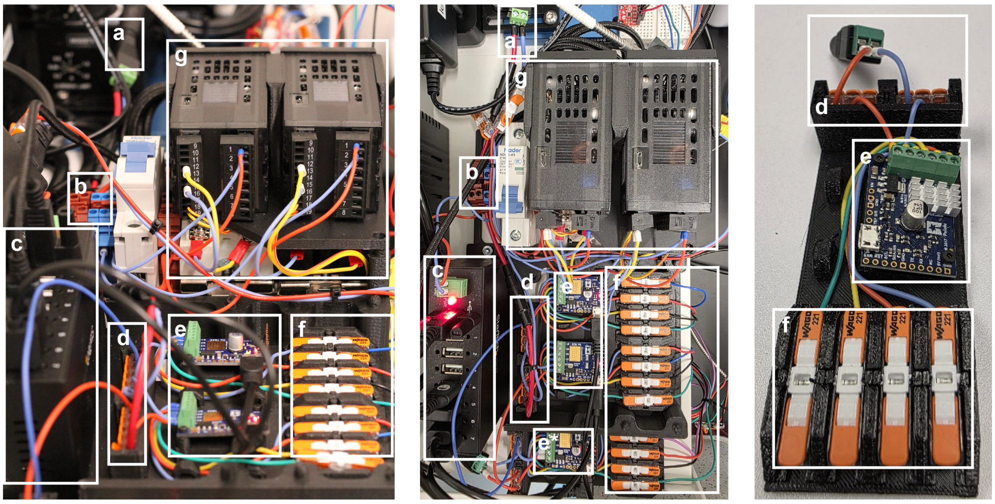


**Fig. S13 Electronics for pump motor and microscope stage control.** **a**, A DC jack to screw terminal breakout couples to a DC power supply jack supplying **b**, a common DIN-mount terminal block shared between motor and heating circuits. An inline fuse provides primary protection to the common power block. **c**, An industrial format USB hub is used to multiplex stepper motor pump controllers and optional microscopy actuators. **d**, Wago terminals couple shared power between motor drivers. **e**, Pololu Tic USB motor controllers are used for pump and **e** microscopy actuator control. **f**, The output pins are broken out in a quick disconnect format coupling to motor drive wires by an array of one-to-one Wago connectors. **g**, The same electronic control module carries the heater control submodule for feedback control of the tissue culture well and gas exchanger heat blocks, comprising thermocouple inputs, PID controllers, solid state relays and relay-switched power outputs fed by the common power supply **b**.

**Table S6** Supplemental Table 6: Bill of materials for heat block manufacturing and control

| **Component or tool description** | **Specification** | **Product identification and/or source** |
|---|---|---|
| Heat block | CNC fabrication | PCBWay fabrication, 5052 aluminum |
| K-type thermocouple | Surface Contact,0.25 mm Diameter, K-Type Sensor Probe with Sticker | PerfectPrime TL0225, Amazon B01NBM7SBK |
| Resistive heaters | Generic 12V 12W 10mmx93mm resistive heater (serpentine trace) | Amazon B09YLRT526 |
| Heater controller | Omega CN401-1445 PID controller | Omega CN401-11445, Newark 34AJ1413 |
| Thermal fuse | Cylinder format 98C thermal cutoff fuse | Cantherm TCO 250VAC 15A 98C (208 F) CYL, Digikey 317-1565-ND |
| Solid state relay | Sensata-Crydom DC relay, >5A >24V rated | Sensata-Crydom 84134750 SSR RELAY SPST-NO 10A 3-60V, Digikey 646-1228-ND or similar |
| Inline fuses | Generic 5x20mm format inline fuse holder and fuse | Amazon B0813Q4S6P or similar |
| Circuit breaker | Single pole DIN rail circuit breaker, 6-10A | ASI-Ez NDB2-63 C10, Digikey 3500-NDB2-63C10-1-ND or similar |
| DIN terminal block for power distribution | Generic | Amazon B07NVV28D9 or similar |
| DIN rail mount for solid state relay | Generic | Amazon B0D8B1J1JZ or similar |
| DIN rail for component mounting | Generic, 200mm length | Amazon B0DGCBMVD1 or similar |
| DC jack to screw terminal adapter | Generic, for 5.5x2.1mm DC jack | Amazon B07LFRDSB7 or similar |

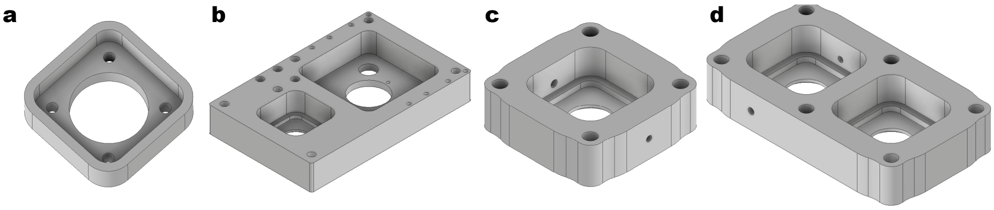


**Fig. S14 Designs for heat blocks.** A joint heat block (above) integrates the heat block for the well device and for the gas exchanger into a monolithic unit. Independent heat blocks (below) for two- and one-well devices allow separate heat control of well device heat blocks and gas exchanger heat blocks.

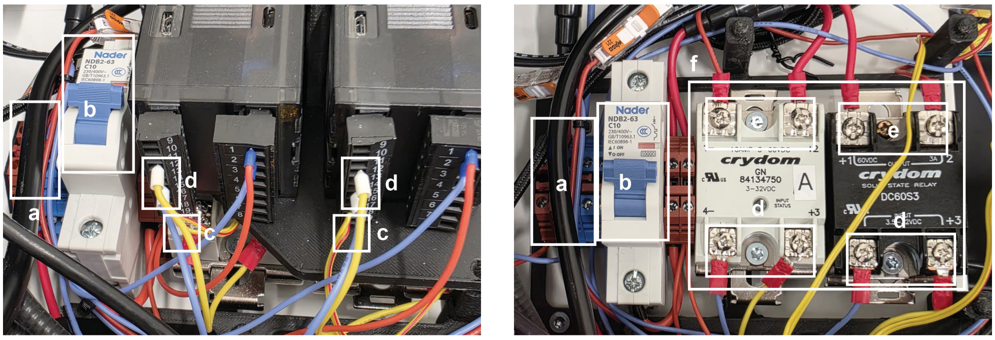


**Fig. S15 Electronics for heat block control.** **a**, A DIN rail mount power distribution blocks and **b**, a heating circuit breaker/manual cutoff are installed on a DIN rail. PID controllers mounted on a carrier are suspended above solid state relays. They receive thermocouple inputs **c** and drive inputs **d** of DC solid state relays **f** switching power supply to resistive heaters **e** on the heat blocks for the tissue culture well and gas exchanger.

**Table S7** Bill of materials for fluidic loop assembly

| **Component or tool description** | **Specification** | **Product identification and/or source** |
|---|---|---|
| Self-sealing disconnect | Silicone swabbable female luer disconnect to male luer lock | Qosina 80213 |
| Gas-impermeable fluidic tubing | Kynar KFlex 1-2, 1/16” ID, 1/8” OD | Eldon James KFLEX1-2 |
| Inflow and outflow check valves | Polycarbonate/silicone 4.2 psi cracking pressure luer lock check valve | Qosina 80136 |
| Male luer to hose barb connector | Generic, polycarbonate luer to 1/16” hose barb | Qosina 11545 or similar |
| Female luer to hose barb connector | Generic, polycarbonate luer to 1/16” hose barb | Qosina 11544 or similar |
| Elbow hose barb connector | Generic, polycarbonate 1/16” elbow hose barb tube-to-tube connector | Colder Products Company HE291 |
| Straight hose barb connector | Generic, polycarbonate 1/16” straight hose barb tube-to-tube connector | Colder Products Company HS291 |
| Tee hose barb connector | Generic, polycarbonate 1/16” elbow hose barb tee connector | Colder Products Company HT291 |
| Wye hose barb connector | Generic, polycarbonate 1/16” elbow hose barb wye connector | Colder Products Company HY291 |
| Effluent line tubing | Tygon S3 E-3603, 1/16” ID, 1/8” OD | Saint Gobain ACF00002 |
| Spin lock male luer to hose barb adapter | Polycarbonate, luer to 1/16” hose barb, spin lock | Qosina 89343 |
| Female to female luer connector, for connecting lines and priming system before inserting well | Generic, polypropylene or polycarbonate female to female luer lock coupler | Qosina 61724 or similar |

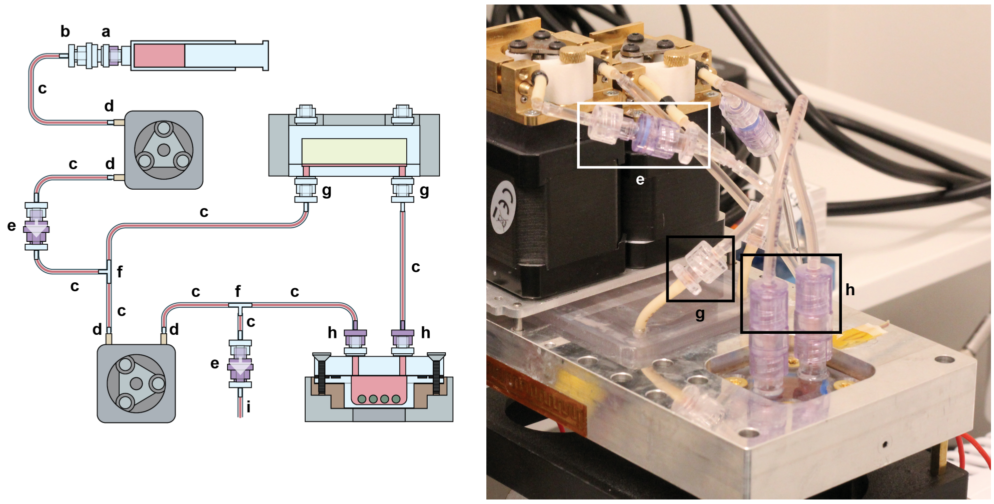


**Fig. S16 Assembly of Fluidic Loop.** **a**, a swabbable female luer self-sealing disconnect forms an aseptic access port for the supply syringe (for sterilant, PBS wash, media) **b**, this is coupled to the inflow line via a female to male luer hose barb adapter. **c**, gas impermeable Kynar KFlex tubing is used throughout the loop, in the configuration pictured. **d**, elbow or straight hose barb tube-to-tube connectors couple tubing to peristaltic pump tubing. **e**, a check valve apparatus is formed by a male hose barb adapter, luer check valve and female hose barb adapter. **f**, inflow and outflow lines access the loop via tee or wye hose barb connectors. Generally we use a tee for inflow and a wye for outflow but they are functionally equivalent. **g**, we use a combination of male and female luer to barb connectors to connect the loop to the inflow and outflow tubing of the gas exchanger. This could equivalently use a tube-to-tube hose barb connector. **h**, to allow installation of the well without twisting of the lines, (since the well lid ports are rotationally constrained) we use a spin-lock rather than simple luer-lock male hose barb to luer adapter in order to couple to the female swabbable quick disconnect ports of the well.

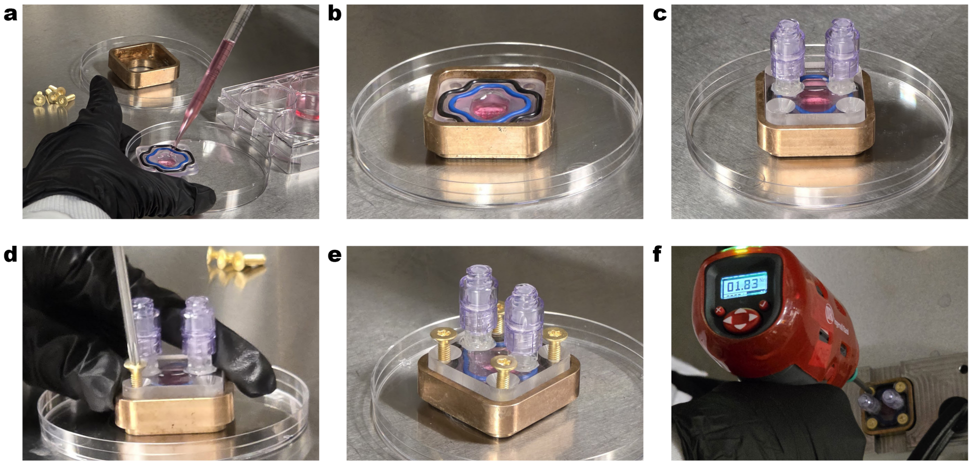

**Fig. S17 Procedure for loading of well with organoids and insertion of screws. a**, Organoids are loaded into the device by pipetting them into the central cavity using a serological or wide-bore pipette tip. **b**, The well is filled to approximately 75% of its volume with media. After sealing, excess air will be removed using the pump's priming routine. Overfilling is avoided, as it could cause bulk overflow beyond the inner gasket or draw fluid past the gasket during sealing. **c**, The well lid is loosely placed over the gaskets, guided by the walls of the inner heat block/well sealing brace. **d**,**e** Brass screws are inserted into the four corner holes and tightened to finger tightness using a T20 Torx L-key, sanitized with Oxivir wipes or autoclaved. At this point, the well is sealed, but the loose seal cannot withstand applied fluidic pressure. **f**, For complete sealing, a CNC milled tightening brace clamped to a table stabilizes the well, and a handheld automated torque screwdriver is used to tighten the four screws to a torque specification of 1.8 Nm.

**Table S8** Bill of materials for well loading and assembly

| **Component or tool description** | **Specification** | **Product identification and/or source** |
|---|---|---|
| Brass clamping brace/heat block component | CNC fabrication | C360 brass, PCBWay |
| Brass sealing screws | T20 Torx drive brass flat head M4 x 14mm screw | Custom fabrication, Shenzhen Shishitong Metal Products Co. Ltd. |
| Autoclave bag | 3.5" x 5.25" autoclave sterilization pouch | PlastCare USA 3.5" x 5.25", Amazon B07K1LY8KL |
| Torx L-key | Chrome plated T20 Torx L-key | Generic eg uxcell, Amazon B0983VLD2Z or similar |
| Tightening fixture | CNC Fabrication | 5052 Aluminum, PCBWay |
| Torque screwdriver | Ingersoll Rand QX Precision Torque Screwdriver | Ingersoll Rand QXFD2PT004PQ04 |
| Ethanol wipe | Coviden sterile alcohol prep pad | Coviden 5750, Amazon B00KJ6U8NE or similar |
| Oxivir wipe | Diversey Oxivir Tb disinfectant wipe | Diversey 5627427, Cole Parmer EW-78901-02 |
| Sanitizing spray | 70% ethanol or isopropanol solution | TexWipe TX3273 Sterile 70% Isopropyl Alcohol Solution or similar |

**Table S9** Bill of Materials for pH measurement apparatus

| **Component** | **Specification** | **Product identification and/or source** |
|---|---|---|
| pH cuvette body | CNC milled fabrication | 3/8" polycarbonate (McMaster Carr 8574K311), CNC milled on Carbide3D Nomad3 desktop mill |
| pH cuvette window | CNC milled fabrication | 1/8" polycarbonate (McMaster Carr 8574K26), CNC milled on Carbide3D Nomad3 desktop mill |
| Alternative pH cuvette | 3D printed fabrication (Asiga Ultra 32) | 3D printed from Detax 04095 Freeprint Ortho 385 Resin, file asiga pH cell.ipt |
| pH shroud body | Fused deposition modeling (FDM) 3D printed fabrication | 3D printed from PLA filament |
| pH shroud lid | FDM 3D printed fabrication | 3D printed from PLA filament |
| Color sensor | Adafruit AS7262 6-Channel Visible Light / Color Sensor Breakout | Adafruit 3779 |
| Screw terminal block connectors for semi-permanent wiring of sensor to Arduino | Generic 2.54 mm pitch screw terminal blocks, through-hole solderable | Amazon B09F6TC7RP or similar |
| Microcontroller interface | Arduino Uno Rev 3 | Arduino A000066 |
| Sensor to microcontroller wiring | Generic 24 AWG wire, multi color kit | Amazon B07G2BWBX8 or similar |
| Wire termination ferrules | Generic 24 AWG wire crimp ferrules | Amazon B07Z5V8818 or similar |
| Screw terminal breakout for semi-permanent wiring of Arduino to sensor | Generic Arduino Uno format screw terminal breakout | Amazon B07HF2DD7T or similar |
| White light reflecting tape | Vinyl electrical tape, white | NSI WW-722-GY |
| Black light blocking tape | Vinyl electrical tape, black | 3M 700 SKU CBG-NAWUS1596 |
| Double sided tape | Foam VHB tape | Gorilla 6055002 heavy duty mounting tape |
| Fluidic connector tubing | 1/16" ID x 1/8" OD x 1/32" Wall PharMed® Tubing | US Plastic Corp 57739 |
| Adhesive for tubing and window | Loctite 4311 | RS Hughes 079340-00003 |
| Primer for tubing | Loctite 7011 | RS Hughes 079340-19886 |

#### Supplemental Discussion

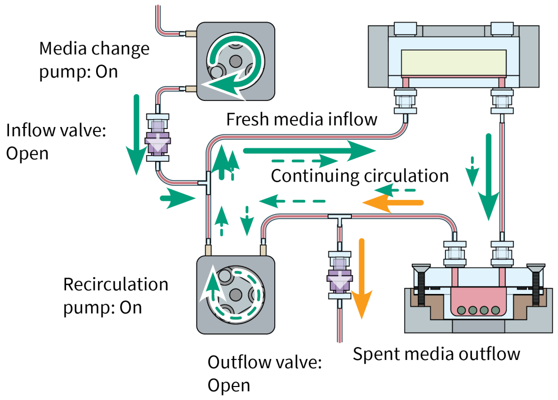


**Fig. S18 Continuous media exchange.** The apparatus described in this paper also allows for a method of media exchange not achievable in drain-and-fill or expansion tank based apparatuses.

Since the sealed, constant-volume loop passively matches media outflow and inflow, not only does automation allow frequent media changes, it is in fact possible to perform continuous "trickle" media replacement as opposed to the conventional discrete time-point media changes commonly used in manual and automated media exchange.

This approach may more closely mimic in vivo physiology in which extracellular fluid constituents are, at the whole-body level, replaced by continuous processes of ingestion, metabolism and biosynthesis and removed by continuous processes of metabolism, filtration and excretion.

This type of media change can be achieved by setting an inflow rate $R$ much lower than the circulation rate but continually running the media change pump. This will produce equal outflow from the loop at rate $R$. The well flow rate may be preserved by setting the recirculation rate to its media change-free rate minus $R$. Then a slow replacement of media, potentially on the order of circulating volume replaced over days, can be achieved without at any point halting circulation across the well. This media exchange will be dilutive due to mixing within the well and/or other regions of turbulent flow in the system.

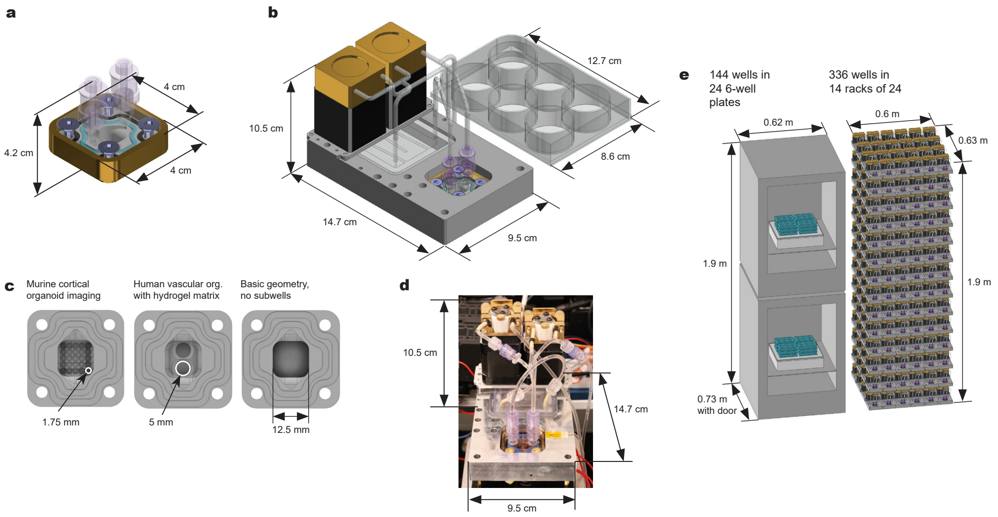


**Fig. S19 Dimensions and scaling.** **a**, Schematic showing dimensions of current design of organoid culture well. This unit is self-contained due to sealing ports and may be transported or stored with organoids independently of the fluidic apparatus. **b**, Schematic showing dimensions of the current design of the assembled fluidic apparatus (without integrated sensors), with comparison to a standard well plate. **c**, Schematic showing internal dimensions of selected organoid culture well geometries. **d**, Photograph showing dimensions as in **b**. **e**, Comparison of the volume occupied by our system versus by conventional shaker based in-incubator organoid culture. We illustrate the space occupied by shaker organoid culture in incubators, noting that typically only 12 plates can fit on one standard CO2 incubator shaker to prevent spillage, and only one shaker can fit in an incubator due to cabling and vibration limits. By comparing our device's dimensions to a pair of stacked incubators in our lab, we show how many replicates of our system could be stored in a rack format within the same volume. The vertical arrangement assumes a server rackmount format with 3U spacing. In this case, we are able to pack 336 wells of organoids in the space normally occupied by 144 wells of organoids in a traditional well plate shaker format. This uses our current implementation of a compact incubator-free culture system, which we have not optimized for a rack format. Optimizing positioning of components would lead to higher density of wells in a rackmount format using our system.

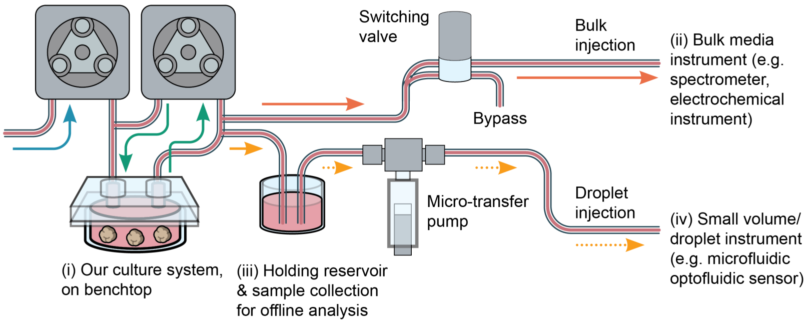


**Fig. S20 Integration with analytical instruments.** Automated online effluent analysis of conditioned media can be achieved by colocating our sealed culture system with analytical instruments on a benchtop **(i)**. For instruments that can accept high-flow effluent directly, an automated switching valve diverts the effluent to the instrument's inlet **(ii)**. For instruments needing low sample volumes, (e.g. droplet based microfluidics) **(iv)**, an expansion reservoir on the effluent line **(iii)** can be used with a microinjection transfer pump for precise volume delivery. The effluent reservoir **(iii)** can also collect fractions for offline analyses.

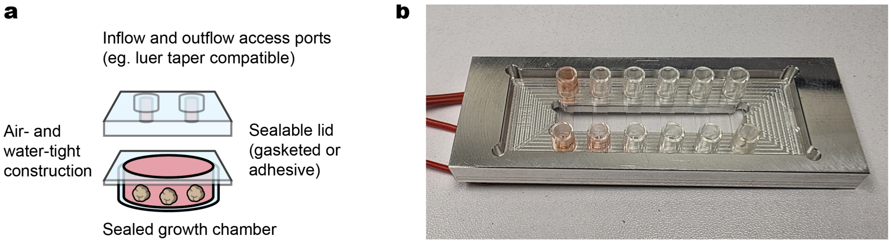


**Fig. S21 Compatibility of our system with alternative tissue culture modules.** The modularity of our apparatus allows for other custom or commercial bioreactor and/or culture wells to be used as well. **a**, Design requirements for a tissue culture device to be coupled to our system. Our system is compatible with any device which can be sealed to be air- and water-tight (e.g. via a clamped gasketed lid, a hermetic plastic-on-plastic seal or an adhesive seal) and which possesses inflow and outflow access ports. Preferably, these access ports will be luer taper format to allow direct use of the fluidic components (e.g. luer swabbable sealed quick disconnects) however, since other fluidic coupling formats (e.g. micro-luer, $\frac{1}{4}$-28 UNF chromatography fitting, barb coupling) are interoperable via adapters, this is not strictly necessary. **b**, We have constructed a microscope slide-format heat block for use of the ibidi µ-Slide VI 0.4 culture device (pictured) in our system. This device consists of six pairs of inflow and outflow access ports serving six independent growth chambers, all sealed by adhesive after seeding of tissue into the growth chamber.
